# Cortex-Wide Substrates for Body Schema and Action Awareness

**DOI:** 10.1101/2025.09.23.678085

**Authors:** Kyle S. Severson, Jinghao Lu, Wenxi Xiao, He Jiang, Ting Lou, Kian A. Caplan, Manuel S. Levy, Tianqing Li, Timothy W. Dunn, Fan Wang

## Abstract

The concept of the body schema originated from studies of patients with cortical lesions who were unable to localize their body parts in space or perceive passive movements (Head & Holmes, 1911). Current consensus defines the body schema as the brain’s internal representation of the body configuration in space used for both embodied perception and voluntary action (de Vignemont, 2010; Berlucchi & Aglioti, 2010; Vallar et al., 2025). Despite extensive research, how the cortex efficiently encodes the vast repertoire of body postures to support natural behavior remains poorly understood. Here, we reveal the representational format underlying the body schema by combining large-scale electrophysiological recordings with full-body 3D tracking and joint angle computation in freely behaving mice. Across cortical regions, neurons encoded diverse postural features in distinct reference frames: posterior parietal cortex (PPC) populations preferentially encoded body midline-referenced features, while sensorimotor cortices utilized a gravity-referenced frame. Strikingly, during performance of different stereotyped actions, distinct cortical ensembles were selectively activated. These action ensembles fired in phase-locked sequences that tiled the full action cycle, providing a neural substrate for action awareness. A network model recapitulated these dynamics and identified low-dimensional action subspaces within population activity. Together, these findings reveal that the cortex organizes the body schema around actions, compressing high-dimensional kinematics into an efficient, body-midline-and gravity-anchored, action-aware neural code.

## Introduction

In the early 20th century, Head and Holmes (1911) completed a study on patients with cortical lesions and discovered that the most common deficits involved an inability to localize body parts in space. Patients also frequently failed to perceive passive movements, for example, when an experimenter moved the patient’s limb without their volition. These observations bore the concept of the “body schema”: the internal representation of the body’s posture in three-dimensional (3D) space. Subsequent research showed that various cortical lesions disrupted body schema and led to numerous neurological symptoms, including impaired awareness of movement and delusional misattribution of body ownership (Bermúdez, 1995; Maravita et al., 2003; Haggard & Wolpert, 2004). Despite its central role in spatial body perception, embodiment, and action, the neural basis of the body schema, specifically its representational format in the brain, remains poorly understood.

Understanding the neural basis of the body schema requires addressing three fundamental questions. The first concerns dimensionality. In vertebrates, postural changes occur at the joints, and mammalian bodies are composed of many joints, with most joints allowing multiple degrees of freedom. This results in a vast, high-dimensional posture space, comprising all possible combinations of joint angles. Proprioceptive afferents innervating muscles, tendons, and joints encode this high-dimensional joint space and transmit detailed, local proprioceptive information to the cortex (Tsay et al., 2016). Does the brain maintain this high-dimensional representation to construct the body schema, or does it encode a more efficient, low-dimensional format?

The second question concerns the reference frame of posture representations. While the body schema is proposed to use egocentric reference frames (Vallar et al., 1999; Medina & Coslett, 2010), by definition, posture is inherently defined as the configuration of the body relative to gravity (Berthoz, 1997; Mittelstaedt, 1997). For example, standing upright and lying flat on the ground may involve nearly identical joint angles in the limbs and torso but are perceived and function as distinct postures due to their orientation with respect to gravity. Gravity thus provides a stable, external reference frame for the body schema. Sensory systems including the vestibular apparatus, proprioception, and vision together offer reliable cues for inferring the direction of gravity (Angelaki & Cullen, 2008; Chesler et al., 2016). Does the brain integrate these multisensory signals to anchor the body schema in a gravity-centered reference frame?

The third closely related question concerns the format of body schema in the context of dynamic actions. Unlike static postures, actions involve coordinated changes in body configuration across the temporal dimension. The body schema must therefore be continuously updated during movement to track these evolving postural states (Haggard & Wolpert, 2004). Yet, we do not consciously perceive these postural changes on a millisecond timescale. Instead, actions are typically experienced as discrete modes, such as running or squatting. How does the body schema incorporate dynamic postural changes during actions? Answering these fundamental questions is critical for understanding how the brain supports motor control and action awareness.

To begin addressing these questions and investigating the neural basis of the body schema, we examined cortical representations of full-body posture in freely moving mice, which naturally exhibit a rich and diverse repertoire of body configurations during spontaneous behaviors. We combined large-scale extracellular recordings with 3D full-body posture tracking to relate cortical activity with naturalistic postural dynamics. Our primary focus was on the posterior parietal cortex (PPC) and the primary and secondary somatosensory cortices (S1 and S2), given their putative roles in body awareness and proprioceptive integration.

The PPC has been implicated in awareness of the body and voluntary actions. In humans, lesions to PPC can cause asomatognosia (loss of awareness of body parts), hemi-spatial neglect (failure to attend to one side of the body), and deficits in recognizing self-generated movements (MacDonald et al., 2003; Sirigu et al., 2004; Vallar & Ronchi, 2009; Vallar & Calzolari, 2018). Electrical stimulation studies further suggest a causal role, with PPC stimulation evoking subjective intentions to move and illusory movement without overt muscle activity at higher stimulation intensities (Desmurget et al., 2009). While these phenomena in humans involve subjective reports, the underlying representational format can be studied in rodents at the neural population level. In rats, PPC encodes lateralized turning during navigation (Whitlock et al., 2012) and efficiently encodes head and trunk posture during natural behavior (Mimica et al., 2018), though it remains an open question whether and how PPC represents the spatial configuration of the entire body, including the tail and limb appendages.

Primary (S1) and secondary (S2) somatosensory cortices are also strong candidate regions involved in the body schema, particularly given their roles in processing proprioceptive input (Medina & Coslett, 2010). Formation and maintenance of the body schema is thought to depend on multisensory integration, combining proprioceptive, tactile, vestibular, and visual inputs (Maravita et al., 2003; de Vignemont 2010; Preuss et al., 2018). Among these, proprioceptive feedback plays a central role in signaling joint angles and postural changes (Roll et al., 1991; Proske & Gandevia, 2012; Tsay et al., 2016). In anesthetized rodents, muscle- and joint-responsive neurons have been identified predominantly in the agranular cortex (primary motor cortex, M1), proprioceptive areas of S1 (Chapin & Lin, 1984; Lee & Kim, 2012), as well as S2, where neurons respond to tendon afferents spanning the entire limb (Marasco et al., 2017). In non-human primates, proprioceptive responses are also distributed across S1, especially area 3a (Prud’homme & Kalaska, 1994), as are responses in awake rodents, which contribute to perception of the body’s position in space (Alonso et al., 2023).

To determine the respective contributions of cortical regions to the body schema, we performed chronic in vivo recordings from PPC, S1, and S2, as well as M1, primary visual cortex (V1), and hippocampus (HPC) for comparison. Across 18 mice and 340 one-hour sessions, we examined how different cortical regions encode full-body posture and movement features during natural behavior. We show that the mouse cortex compresses high dimensional full-body kinematics into low-dimensional representations anchored to gravity and the body midline, organized temporally by current action identity and phase, providing a neural substrate for an action-aware body schema.

## Results

### Full-body posture tracking and joint angle computations

To investigate how the cortex encodes the body schema during natural behavior, we developed a motion capture system for tracking full-body posture in freely moving mice. Spontaneous behavior was recorded in an enclosed arena surrounded by six synchronized, calibrated cameras, enabling high-precision, non-invasive 3D tracking (Fig. 1a). We defined a comprehensive skeletal model with 26 anatomically grounded keypoints (Extended Data Fig. 1a,b) which were used to train a DANNCE model (Dunn & Marshall et al., 2021) for markerless 3D pose estimation, achieving a mean Euclidean error of 2.38 mm on held-out test frames (Fig. 1b-d). Once the pose estimation model was validated, we predicted keypoints for a total of 122.4 million frames across 18 mice. All 3D keypoint predictions were calibrated to the center of the arena floor, representing continuous x-, y-, and z-coordinate time series for each keypoint (See Supplementary Video 1).

**Figure 1.**
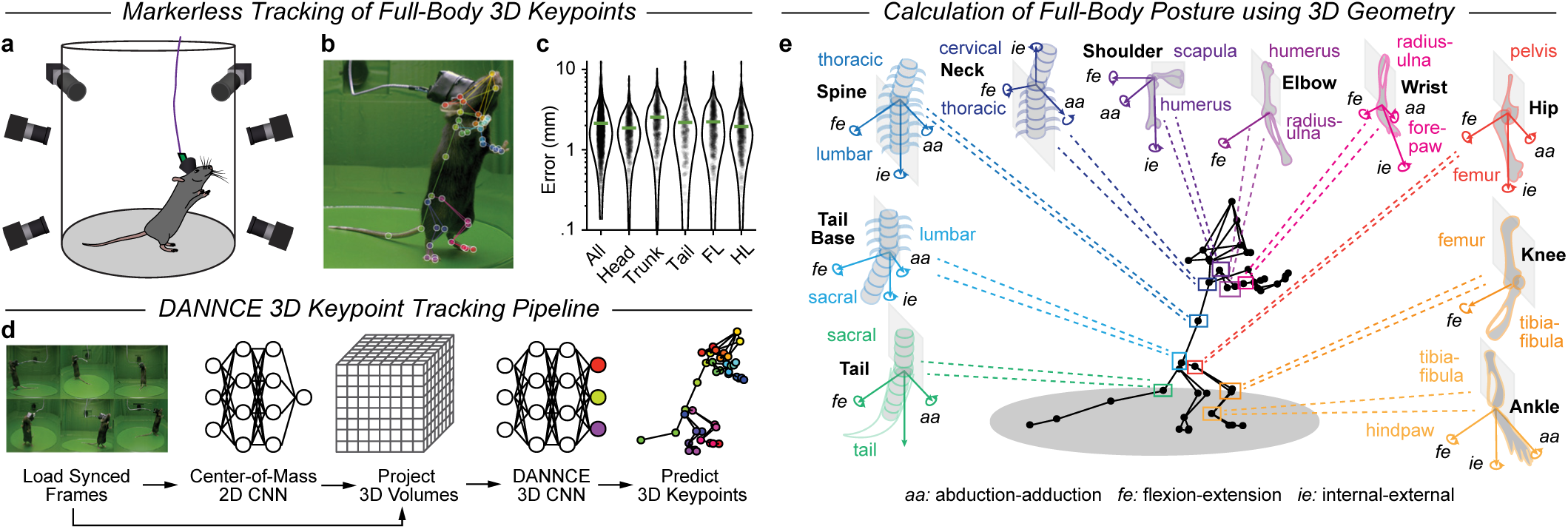
Tracking full-body kinematics with 3D geometric models. **a**, Markerless motion capture setup with six synchronized cameras surrounding a mouse freely behaving in an enclosed arena. **b**, Single-camera frame with DANNCE keypoint predictions (colored circles) connected by links (colored lines). The mouse was implanted with a tethered 64-channel microdrive for simultaneous electrophysiology. **c**, Euclidean distance between human-labeled and DANNCE-predicted 3D positions across validation frames (n=60 frames, 12 mice); green lines, median error for each body part; mean error 2.38 mm. FL: forelimb; HL: hindlimb. **d**, DANNCE markerless pose estimation applies 2D CNNs to multi-view frames to triangulate the 3D center-of-mass (COM), projects the video frames into a COM-centered volumetric grid via projective geometry, and uses a 3D CNN to predict 3D keypoint positions. **e**, Anatomical skeletal model estimating joint angles via 3D trigonometry from the rotation of the distal body segment relative to each proximal segment.

To derive interpretable measures of head and body orientation, we grouped these keypoints into 17 anatomically defined body segments spanning the head, trunk, limbs, and tail (Extended Data Fig. 1c). To compute the 3D positions and orientations of these body segments, we defined each segment as a plane comprised of three adjacent keypoints (e.g., snout and ears for the head) and decomposed world-centered segment rotations into Euler angles (Extended Data Fig. 1e). Yaw represents allocentric (world-centered) heading, while pitch and roll capture segment orientation relative to gravity.

Because the size and shape of body segments in adult mice remain constant during movement (Extended Data Fig. 1d), postural changes arise from joint rotations driven by muscle activity (Monsees et al., 2022). Rotations at the joints and head thus serve as biomechanical correlates of proprioceptive and vestibular input, key components from which the brain constructs the body schema. We constructed a joint-centered geometric model of the skeleton by defining 16 joints connecting adjacent body segments (Extended Data Fig. 1f-h). Each joint was assigned up to three rotational degrees of freedom to parameterize full-body posture (Fig. 1e). Ball-and-socket joints (e.g., shoulder, hip) allow flexion-extension (*fe*), abduction-adduction (*aa*), and internal-external rotation (*ie*), while hinge joints (e.g., elbow, knee) support only flexion-extension. This joint-centered geometric skeletal model provides the basis to compress the posture parameter space into 40 interpretable dimensions (38 joint angles, head pitch and roll) that capture the body configuration and its orientation with respect to gravity at each time point. When enriched with derived postural dynamic features such as joint velocities, our framework forms the foundation to link cortical activity to detailed full-body kinematics in freely behaving mice.

### PPC preferentially encodes postural angles of midline body parts

To examine how cortical neurons encode body posture, we first performed extracellular recordings from the medial PPC by implanting chronic tetrode microdrives posterior to the S1 trunk region (Fig. 2a,b; N=4 mice). In total, we isolated 966 single units sampled across all cortical depths (see Extended Data Fig. 2a,c), while simultaneously acquiring behavioral video for automated full-body tracking (61 sessions, totaling 3,660 minutes). Inspecting these data, we observed that neural activity was modulated by changes in posture, including shifts in body height, head pitch, and specific joint angles (e.g., Fig. 2c,d). To systematically quantify posture tuning, we computed tuning curves for every neuron to 58 posture features, including joint angles, body segment orientations, and height of different body parts, and compared their relative tuning strength using variance explained (*R*²) as a metric (Extended Data Fig. 3a,b,e; Methods). Among the 738 regular-spiking (RS) PPC neurons (identified by waveform analysis; see Extended Data Fig. 2d-f), 52.3% (386/738) were tuned to one or more posture features. A heatmap of normalized tuning curves across the posture-tuned population revealed that most neurons preferred extreme angles (Fig. 2e), consistent with efficient coding of posture angles observed in PPC (Mimica et al., 2018).

**Figure 2.**
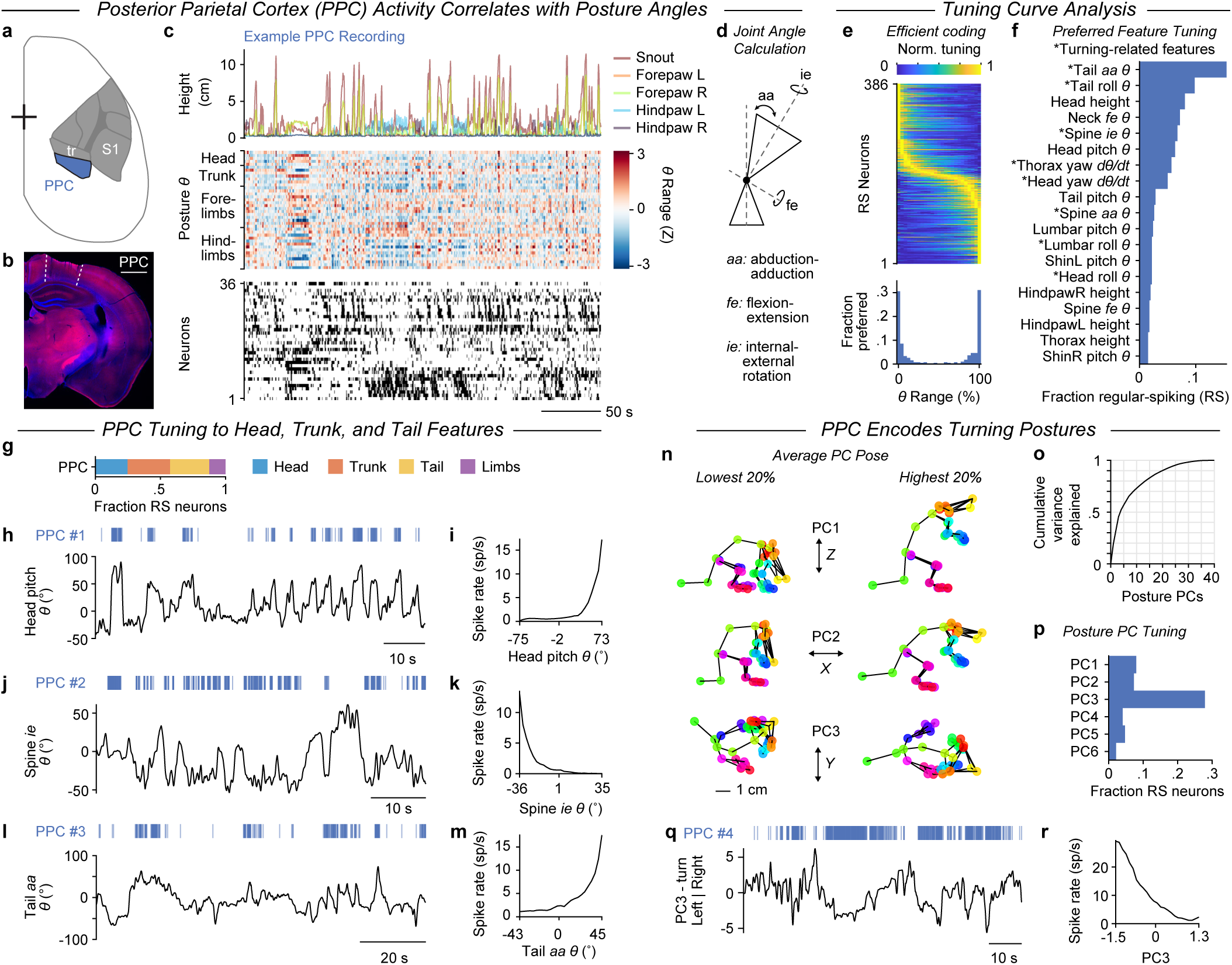
PPC encodes turning postures through preferential tuning to midline trunk angles. **a**, Dorsal view of mouse right hemisphere with medial PPC outlined. Tetrode microdrives were chronically implanted in right PPC, posterior to the empirically mapped S1 trunk (tr) region identified via intrinsic optical signal imaging. **b**, Cortical section of the right hemisphere after tetrode removal, showing the target-ed PPC region. Scale bar, 1 mm. **c**, Time-aligned example traces. Top, keypoint height (cm) of distal body parts. Middle, Z-scored time series of 40 posture angles. Bottom, spike raster of 36 simultaneously record-ed PPC neurons, sorted by Rastermap; grayscale firing rates were Z-scored and saturated (range: 0–0.1) to highlight activations. **d**, Schematic of abduction-adduction (aa), flexion-extension (fe), and internal-ex-ternal (ie) rotation angles measured at the joint (black circle). **e**, Top, normalized tuning curves for the best-tuned posture variable across all posture-tuned regular-spiking (RS) neurons in PPC, sorted by preferred angle bin. Bottom, histogram of preferred angle percentile bins. Preference toward the extremes suggests efficient coding. **f**, Histogram of best-tuned features for all kinematics-tuned RS neurons (n=459), sorted by frequency. *, turning-related features. **g**, Distribution of best-tuned kinematic features grouped by body part for the same RS neurons in (f). **h-i**, Example spike raster (h) and tuning curve (i) for neuron #1 aligned to head pitch. **j-k**, Same for neuron #2 aligned to spine ie rotation. **l-m**, Same for neuron #3 aligned to tail aa joint angle. **n**, Average poses from lowest and highest 20th-percentile bins of first three Posture PCs. PC1-3 correspond to coarse postural adjustments in *Z-*, *X-*, and *Y*-axes, respectively (see Extended Data Fig. 4). **o**, Cumulative variance in posture angles explained by Posture PCs. **p**, Histogram of tuning preference to top six Posture PCs for all RS neurons (n=738); PC3 captured turning postures. **q-r**, Spike raster (q) and tuning curve (r) for example neuron #4 aligned to PC3.

To understand which posture features were most strongly represented in PPC, we examined the fraction of neurons best-tuned to each individual postural feature. Consistent with prior work (Mimica et al., 2018), we observed prevalent tuning to head, neck, and trunk angles (Fig. 2f). Extending these findings, our full-body tracking revealed that relatively few PPC neurons were tuned to limb-related features (58/459, 12.6%), whereas a surprisingly large fraction was tuned to tail angles (139/459, 30.3%; Fig. 2g), a body part not tracked in earlier studies. In total, 87.4% (401/459) of posture-tuned PPC neurons encoded postural angles of midline body parts, such as the head, neck, spine, and tail (3 example neurons in Fig. 2h-m). Across the PPC population, we found an overrepresentation of posture features reflecting left-right axial trunk rotation, including tail *aa*, tail roll, spine *aa,* and spine *ie* angles, as well as yaw angular velocity of the head or thorax (Fig. 2f). These observations suggest PPC preferentially represents postural deviations from the body midline, particularly those associated with turning.

To reveal low-dimensional postural features, we applied principal component analysis (PCA) to identify the “eigenposes” that capture the dominant axes of the body configuration (see Methods). The first six Posture PCs explained 61% of the variance in full-body posture angles (Fig. 2o; Methods). In order to interpret these eigenposes, we visualized the extreme bins of each Posture PC as averaged 3D poses, revealing that the top three Posture PCs represented orthogonal posture dimensions: PC1 captured elevation changes, PC2 captured forward-backward pitch, and PC3 captured left-right turning of the main body axis including the tail. Together, these Posture PCs describe coarse postural adjustments of the center-of-mass relative to gravity or the body’s midline (Fig. 2n, see also Extended Data Fig. 4).

We hypothesize that these low-dimensional features could be represented in the brain. To test this hypothesis, we asked how PPC neurons were tuned to each Posture PC. Notably, a substantial fraction of RS neurons (27.9%, 206/738) were preferentially tuned to PC3 (Fig. 2p), confirming a population-level preference for left-right turning, consistent with our tuning analysis (Fig. 2f) and prior findings in rats (Whitlock et al., 2012). We observed PPC neurons tuned to PC3, showing activity selective for leftward or rightward turns (e.g., Fig. 2q,r).

Together, these results reveal that PPC neurons predominantly encode head, trunk, and tail angles associated with left-right turning. Deviations from the body’s midline emerged as a central organizing axis for the posture representation in mouse PPC, suggesting that PPC emphasizes a midline-referenced body schema that supports lateral stabilization and turning. Although left-right turning can covary with body roll (a gravity-related component, see Extended Data Fig. 4b,c), roll angles were minimal under our recording conditions. Because posture is defined relative to gravity, we next asked whether a broader parietal network anchors the body schema to a gravity-centered reference frame.

### Regular-spiking neurons in sensorimotor cortices represent posture in a gravicentric reference frame

We expanded our recordings to sensorimotor cortices (SMC) and compared posture tuning across cortical areas. Specifically, we targeted S1 (n=3,928 neurons from 6 mice; see Extended Data Fig. 2b,c), including the proprioceptive subregions S1pgz, S1dz, and S1tz (Fig. 3a), as well as S2 (n=941 neurons from 2 mice). For comparison, we conducted additional recordings in M1 (n=688 neurons from 3 mice), primary visual cortex (V1, n=654 neurons from 2 mice), and hippocampus (HPC, n=1,027 neurons from 5 mice). Tuning analysis revealed prominent posture tuning in RS neurons across SMC (M1: 194/454 neurons, 42.7%; S1: 1,166/2,990, 39.0%) and PPC (386/738, 52.3%). In contrast, posture tuning was relatively sparse in RS neurons within S2 (140/689, 20.3%), V1 (86/530, 16.2%), and HPC (18/496, 3.6%). We expanded the Posture PC analysis to determine how RS neurons in different regions encoded posture. Unlike PPC, which overrepresented PC3 left-right turning, posture-tuned RS neurons in SMC were most strongly tuned to PC1 (Fig. 3b; M1: 84/454, 18.5%; S1: 648/3,525, 15.9%; S2: 55/689, 7.5%), corresponding to changes in body height relative to the floor (Fig. 3c). A sizeable fraction of RS neurons in S1 also showed preference for PC2 (272/3,525, 7.7%), corresponding to backward-forward leaning.

**Figure 3.**
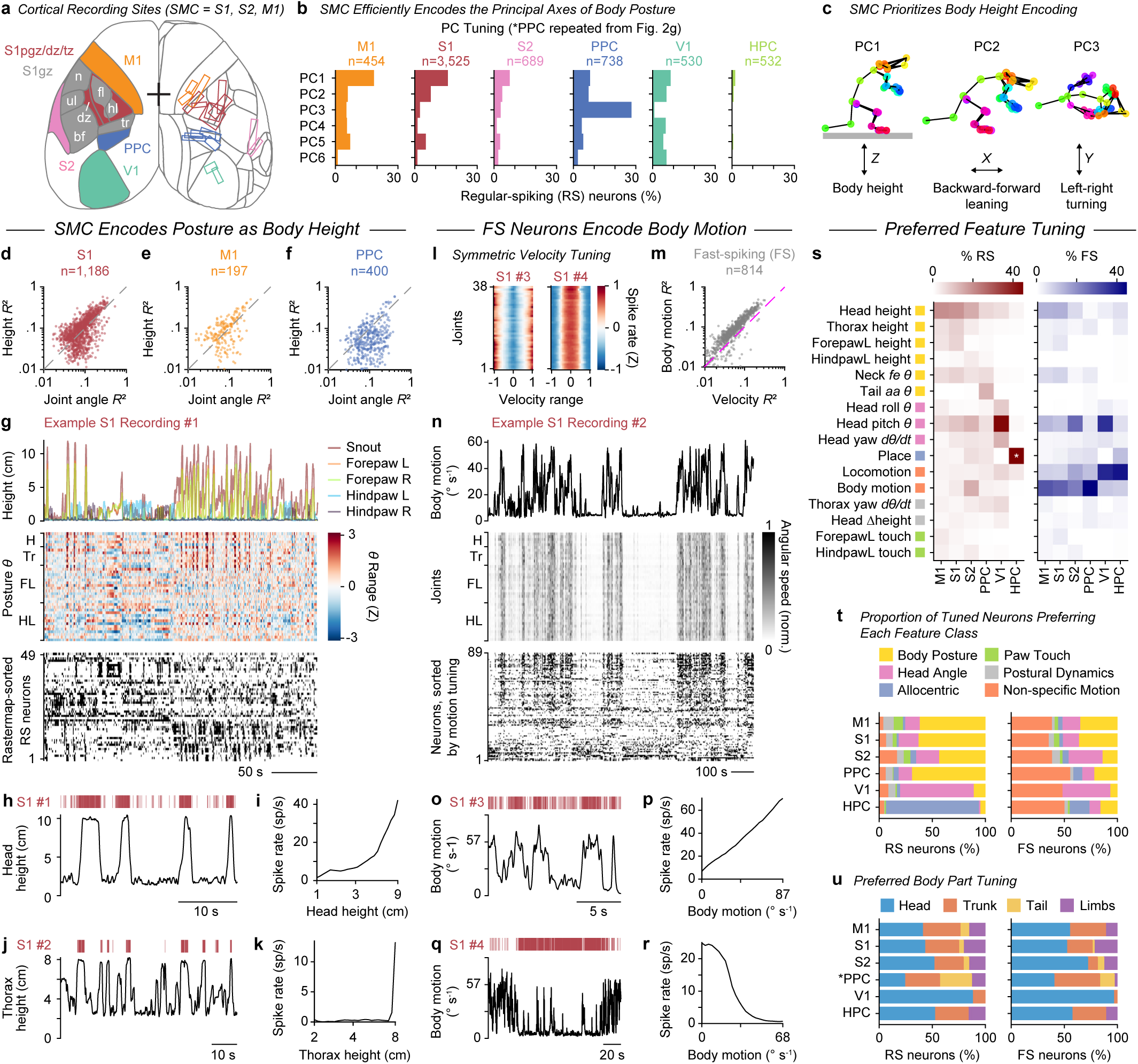
SMC RS neurons encode global posture in a gravicentric reference frame, while FS neu-rons encode non-specific motion. **a,** Dorsal cortex atlas based on the Allen CCF (gray outlines). Black crosshair: bregma (length = 1 mm). Left hemisphere, Targeted regions: M1 (orange), S2 (pink), PPC (blue), V1 (green), and S1 (red) proprio-ceptive zones (pgz, perigranular; dz, dysgranular; tz, S1-M1 transition). Granular S1 (S1gz, dark gray) shows somatotopic zones for snout (sn), barrel field (bf), trunk (tr), hindlimb (hl), and forelimb (fl). Right hemisphere, microdrive locations outlined (n=18 mice, same colors as left). Hippocampus (HPC) was targeted below PPC and posterior S1. **b**, Tuning preference histograms for top six Posture PCs over all RS neurons in each region. **c**, Average poses from first three Posture PCs, representing body height (Z), lean-ing (X), and turning (Y). **d-f**, Scatter plots comparing variance in firing rates explained by the best joint angle tuning model versus best segment height model in S1 (d), M1 (e), and PPC (f) RS neurons. **g**, Time-aligned example traces. Top, keypoint height (cm) of distal body parts, including left (L) and right (R) fore- and hindpaws. Middle, Z-scored posture angles, grouped by body part: head (H), neck/trunk (Tr), forelimbs (FL), and hindlimbs (HL). Bottom, spike raster of 49 simultaneously recorded S1 neurons. **h-i**, Example spike raster (h) and tuning curve (i) for S1 neuron #1 aligned to head height. **j-k**, Same for S1 neuron #2 aligned to thorax height. **l**, Z-scored tuning curves for all joint angular velocities (*dθ/dt*) for exam-ple S1 neurons #3 and #4. **m**, Scatter plot comparing variance in tuned FS neuron firing explained by body motion versus joint angular velocity (see Methods). **n**, Top, example body motion trace, computed from mean joint speed (middle). Bottom, spike raster of 89 simultaneously recorded neurons, sorted by motion tuning slope. **o-p**, Example spike raster (o) and tuning curve (p) for S1 neuron #3 activated by body motion. **q-r**, Same as (o-p) for S1 neuron #4, suppressed by body motion. **s**, Histograms quantifying fraction of RS (left, red) and FS (right, blue) neurons preferring each of the top selected features. * Place for HPC exceeds the color scale range at 84.5%. Colored squares next to the y-axis labels indicate feature class (see legend in t). **t**, Distribution of preferred feature classes for kinematics-tuned RS (left) and FS (right) neurons across cortical regions. **u**, Histograms of tuned neurons with preference for head-, trunk-, tail-, or limb-related features for RS (left) and FS (right) neurons across regions. *, Data from PPC RS neurons in (b), (c), and (u) are repeated from Fig. 2 for comparison.

To directly compare encoding of global postural features in different reference frames, such as body segment height versus local joint configurations, we compared the *R²* of each neuron’s best height feature to that of its best joint angle feature. In both S1 and M1, a greater proportion of RS neurons showed stronger tuning to height relative to the floor (Fig. 3d,e), whereas RS neurons in PPC exhibited the opposite pattern, with stronger tuning to joint angles (Fig. 3f). These properties were clearly evident in population activity, for example in rasters showing widespread modulation by changes in snout, forepaw, and hindpaw heights (Fig. 3g), and in single-neuron traces (Fig. 3h-k). Together, these results indicate that SMC preferentially encodes posture in a gravicentric reference frame, emphasizing global body height and pitch rather than local joint angles.

### Fast-spiking neurons in SMC and PPC robustly encode body motion

In addition to encoding static postures, many cortical neurons were modulated by dynamic posture changes. While analyzing tuning to kinematics (postural features and their temporal derivatives) we found a substantial fraction of FS putative interneurons across M1, S1, S2, and PPC exhibited symmetric tuning to joint angular velocities, regardless of direction and joint identity (Fig. 3l). Thus, rather than encoding the velocity of a specific joint, these neurons were more strongly driven by the average angular speed across all joints, which we define here as “body motion” (Fig. 3m,n). Neurons in SMC and PPC tuned to body motion (RS: 563/4,871, 11.6%; FS: 520/1,652, 31.5%) typically increased their firing rates in proportion to body motion (e.g., Fig. 3n-p; RS: 397/563, 70.5%; FS: 509/520, 97.9%), although a small subset showed the opposite pattern, firing more when body motion was slower (e.g., Fig. 3n,q,r; RS: 166/563, 29.5%; FS: 11/520, 2.1%).

Body motion is distinct from locomotion: while locomotion refers to translocation of the body in world coordinates (i.e., change in X-Y position), body motion refers to movement of body parts relative to the animal’s own body frame (i.e., average joint speed). Locomotion and body motion are correlated during walking but become dissociated during stationary actions, such as self-grooming. When comparing tuning to locomotion versus body motion, more FS neurons in V1 and HPC were tuned to locomotion (Fig. 3s, right), consistent with prior work (Niell & Stryker, 2010; Sun et al., 2018). In contrast, more FS neurons in M1, S1, S2, and PPC showed stronger tuning to body motion compared to locomotion (Fig. 3s, right). Across all cortical regions, a substantial fraction of FS neurons was best tuned to either body motion or locomotion, collectively categorized as “Non-specific Motion” (Fig. 3t, right; M1: 38.3%, 46/120; S1: 35.3.1%, 177/501; S2: 38.5%, 47/122; PPC: 55.7%, 59/106; V1: 48.3%, 29/60; HPC: 50.2%, 113/225). Together, these findings suggest that putative inhibitory interneurons ubiquitously encode non-specific motion, with SMC and PPC specialized for encoding body motion, complementing the posture-selective tuning of putative excitatory neurons in those regions.

### Comparison of kinematic feature tuning across cortex

We next asked how tuning to different categories of kinematic features was distributed across cortical regions. We computed tuning strength for each neuron and each feature (285 total features) and compiled a histogram of preferred feature tuning for RS neurons (Fig. 3s, left). SMC contained large proportions of RS neurons best-tuned to body height features, especially head height. Tuning to forepaw and thorax height was more pronounced in S1 than in S2 or M1. Across SMC, head pitch was also strongly represented, especially in S2. Next, we assigned each neuron into six broader functional categories based on their preferred tuning feature: Body Posture, Head Angles, Allocentric, Paw Touch, Postural Dynamics, and Non-specific Motion (see Extended Data Fig. 3f; Methods). Histograms comparing the distribution of best-tuned features across regions revealed that Body Posture, comprised of joint angles, segment heights, and posture PCs, was the dominant tuning category in RS neurons within M1 (61.7%, 171/277), S1 (62.9%, 1,026/1,631), S2 (43.5%, 111/255), and PPC (69.0%, 368/533; Fig. 3t, left).

Only a small fraction of RS neurons in SMC (6-10%) were best tuned to Postural Dynamics (joint or segment velocities and accelerations; see Extended Data Fig. 3c,d; Methods), suggesting that dynamic features were less emphasized than static postural configurations across cortical RS populations. Few RS neurons in SMC (4-9%) were categorized as encoding Paw Touch features, derived from the proximity of the paws to the floor, walls, or head. In V1, Head Angle features, especially head pitch and head yaw velocity (head turning), were strongly preferred by the RS population (69.3%, 95/137), consistent with encoding the head’s orientation and movement relative to the external world (Mimica et al., 2023; Orlowska-Feuer et al., 2022). As expected, the majority of RS neurons in HPC were Allocentric (84.5%, 251/297), preferring “place” above all other features (Fig. 3t, left).

Across all cortical regions and cell types, head- and trunk-related kinematic features were overrepresented (Fig. 3u). Tail features were more strongly represented in PPC than in M1, S1, or S2, while head-related kinematics were overrepresented in V1. Surprisingly, only 21.5% (304/1,412) of S1 RS neurons showed preferential tuning to limb-related features, despite the fact that we targeted S1 subregions known to receive primary proprioceptive input from the limbs (Chapin & Lin, 1984). These findings suggest that, rather than encoding high-dimensional, primitive proprioceptive inputs, SMC compresses posture into a low-dimensional representation centered on body height relative to the ground and pitch relative to gravity. This gravicentric posture representation, referenced relative to the ground and aligned with the gravitational vector, provides a stable and ethologically relevant reference frame for the body schema. Moreover, through regional and cell-type specialization, the cortex maintains distinct but complementary representations: RS neurons in SMC encode posture as height and pitch, RS neurons in PPC encode midline turning postures, and FS neurons encode non-specific motion. Together, these low-dimensional representations support a body schema distributed across specialized cortical subnetworks.

### Prominent action ensembles in the cortex

We next asked how the body schema incorporates dynamic postural changes during natural behavior. If the brain were to continuously track the instantaneous configuration of the whole body in fine detail, it may be metabolically costly and computationally inefficient. Animals do not move through the world by evaluating each posture as a momentary snapshot. Instead, animals compose spontaneous behaviors from a repertoire of species-typical actions, such as walking, grooming, or rearing, that are stereotypical and repeated routinely throughout their lives (Marshall et al., 2021). For example, during walking, we do not consciously monitor joint angles across each footstep; we simply lean forward and initiate a ‘walking’ sensorimotor program that implicitly guides coordinated movement. We hypothesized that the brain similarly employs an action-level representation that abstracts away the finer details of postural configurations, allowing the body schema to represent postures in a compressed format contingent on behavior context.

To test our hypothesis, we examined how neuronal activity was modulated by actions, defined here as dynamic, stereotyped movement trajectories centered around distinctive full-body postures. In our behavioral arena, mice primarily performed nine common actions: Walk, Rear, Crouch, Sniff, Head Groom, Body Groom, Left or Right Scratch, and Rest (Fig. 4a). We annotated example periods with these categorical labels and used these labels to train a supervised action segmentation model. We implemented an encoder-decoder temporal convolutional network (AS-EDTCN; based on Lea et al., 2017) to classify actions from kinematic data (see Extended Data Fig. 5, Methods). The model achieved high accuracy on held-out subjects (81%) and aligned more closely with human annotations than an unsupervised model (Weinreb et al., 2024; see Extended Data Fig. 5a-c). Sorting neural activity with Rastermap (Stringer et al., 2025) revealed large clusters of neurons that were selectively activated during specific actions, such as Sniff or Groom in S1 (e.g., Fig. 4b) and Rear or Scratch in PPC (e.g., Fig. 4c). These observations suggested that distinct groups of neurons are selectively engaged during specific actions, forming what we term “action ensembles”.

**Figure 4.**
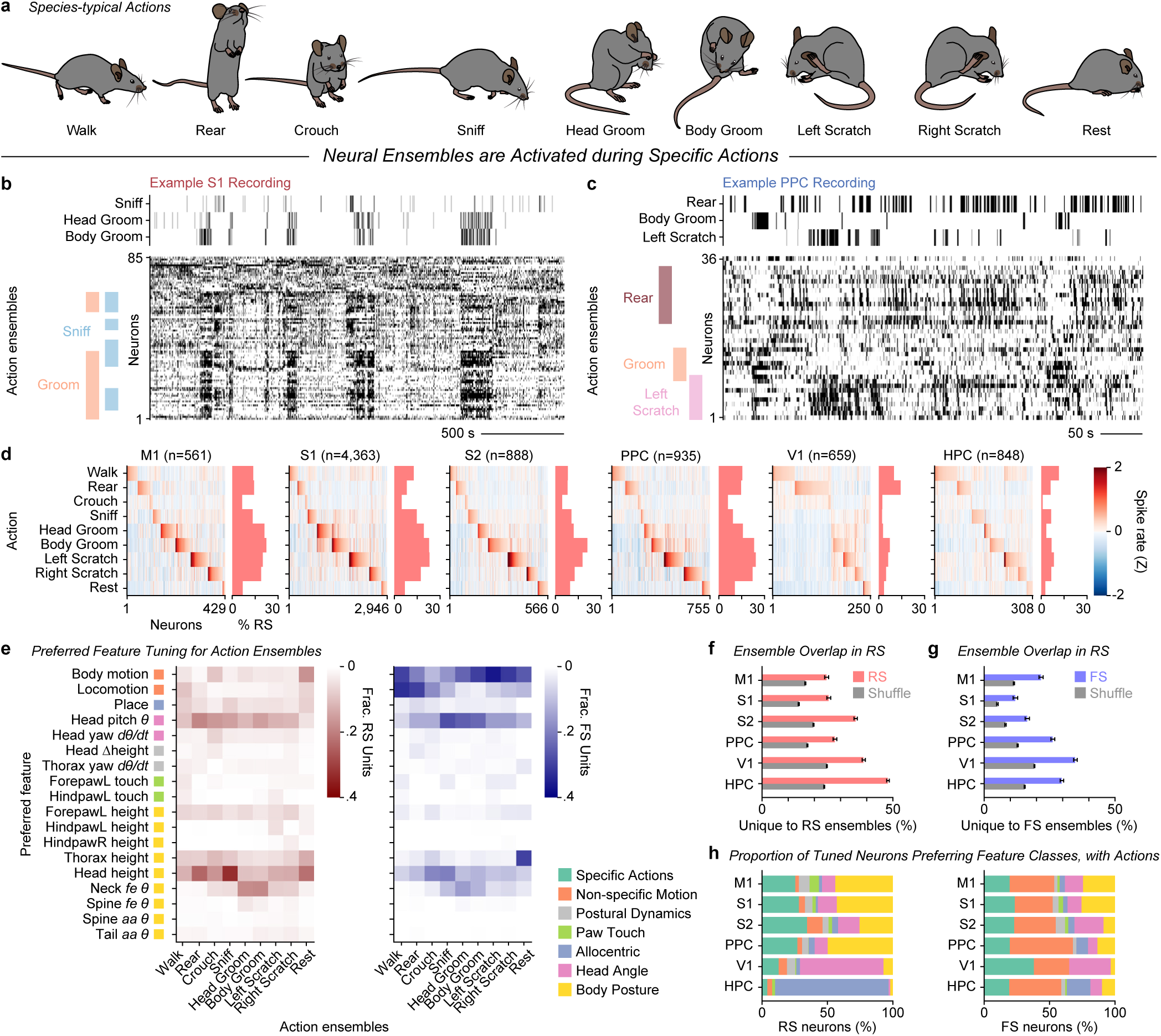
Prominent action ensembles in SMC and PPC. **a**, Nine common species-typical actions observed during spontaneous behavior. **b**, Example snippet from behavioral and S1 recordings in a mouse. Top, ethogram showing occurrence of Sniff, Head Goom, and Body Groom actions classified by AS-EDTCN. Bottom, Spike raster aligned in time, sorted by Rastermap. For illustrative purposes, clusters of neurons aligned to repeated actions were qualitatively labeled as action ensembles. **c**, Same as (b), but for PPC activity aligned to Rear, Body Groom, and Left Scratch actions. **d**, Heatmaps showing mean Z-scored spike rate of action-activated RS neurons (columns) for each action (rows). Histograms showing percentage of regular-spiking (RS) neurons in each region belonging to each (non-exclusive) action ensemble. **e**, Heatmaps showing the fraction of RS neurons (left) and FS neurons (right) within each action ensemble (columns) that prefer each of the top-ranked features (rows). Colored squares adjacent to feature labels indicate feature class (legend shown to the right). **f-g**, Mean percentage of unique neurons per ensemble not shared with other ensembles, compared to unique neurons drawn from shuffled populations (gray) with matched ensemble sizes. Red: RS neurons; blue: Fast-spiking (FS) neurons. Error bars: 95% CI. **h**, Distribution of tuned neurons preferring feature classes for RS (left) and FS (right) neurons across cortical regions. Same format as Fig. 3t, but recalculated with the addition of the Specific Actions feature class. Same feature class legend as in (e).

Building on these observations, we next sought to identify neurons whose firing rates were significantly modulated by individual actions. To identify action ensembles, we used a simple threshold on the mean Z-scored action modulation for each neuron (Fig. 4d; see Methods). This straightforward classification approach revealed prominent action ensembles across cortical areas, especially for Rear, Groom, and Scratch. Remarkably, SMC and PPC contained large proportions of action-activated neurons (PPC: 80.7%, 755/935 RS neurons; M1: 76.5%, 429/561; S1: 67.5%, 2,946/4,363; S2: 63.8%, 566/888), whereas V1 and HPC contained smaller fractions (V1: 37.9%, 250/659 RS neurons; HPC: 36.3%, 308/848) with weaker action modulation (Fig. 4d; see Extended Data Fig. 7a for FS neurons). We also observed action-suppressed ensembles (Extended Data Fig. 6). Moreover, neurons within each action ensemble exhibited kinematic tuning profiles that aligned with the defining features of the corresponding action (Fig. 4e). For example, more RS and FS neurons in the Walk ensemble were tuned to locomotion speed, more RS neurons in the Rear ensemble were selective for head pitch and head height, and a greater number of RS neurons in the Left and Right Scratch ensembles were tuned to hindpaw height.

Having observed prominent neural ensembles representing each of these nine common actions, we next asked whether individual neurons were dedicated to a single action or contributed to multiple action ensembles. By our definition, action ensembles were not exclusive, so neurons could participate in more than one ensemble. To examine this overlap, we computed a matrix of shared ensemble membership across all action pairs (see Extended Data Fig. 7b). Overlap was most pronounced among Head Groom, Body Groom, and Scratch ensembles, likely reflecting shared postures and movement trajectories. On average, 25-50% of RS neurons within each action ensemble were exclusive to that ensemble across cortical areas (Fig. 4f). A significant portion of FS neurons were exclusive to action ensembles, though more promiscuous than RS neurons (Fig. 4g). These findings suggest that stereotyped actions are represented by distinct, but partially overlapping ensembles, likely involving recurrent subnetworks of excitatory and inhibitory neurons.

We next asked how this Specific Actions feature class explained neural variance compared to other kinematic feature classes. Incorporating Specific Actions into the preferred feature analysis revealed that sizeable fractions of tuned RS neurons in SMC and PPC now showed strongest tuning to discrete action categories compared to continuous kinematic features in other classes (Fig. 4h, left; M1: 25.9%, 75/290; S1: 27.8%, 1,718/3,525; S2: 34.4%, 97/282; PPC: 27.0%, 148/548). Nonetheless, the largest fraction of RS neurons remained tuned to Body Posture. The majority of FS neurons across cortical areas continued to be best tuned to Non-specific Motion, although a large portion now preferred Specific Actions (Fig. 4h, right). Together, these results demonstrate that cortical populations form distinct action ensembles, with RS neurons preferentially encoding specific actions and postures and FS neurons contributing to broader, motion-related modulation across actions. These prominent action ensembles may serve as a neural substrate for action awareness, an internal representation of which action is being executed.

### Action ensembles encode phase as neural sequences

Actions unfold as characteristic sequences of posture transitions, each with their own unique physical demands across the stereotypical trajectory. For example, a Rear is typically a single-cycle action in which the animal lifts its forepaws off the ground, rapidly extends its hindlimbs to reach peak height, then descends back into a crouched or prone position (Fig. 5a). In contrast, Head Groom consists of repeated cycles in which the mouse extends its forepaws from behind the ears toward the snout, then retracts them before initiating the next cycle (Fig. 5b; see Extended Data Fig. 8a-b). Actions like Rear, Head Groom, and Walk are therefore inherently phasic, consisting of stereotypical sequences of postures, whereas Rest and Crouch involve relatively static postures. Critically, phasic actions traverse similar postures on the forward and return phases, but with opposite velocity directions. Thus, a neuron locked to phase is not simply encoding a static posture, but rather a unique state within the circular action trajectory.

**Figure 5.**
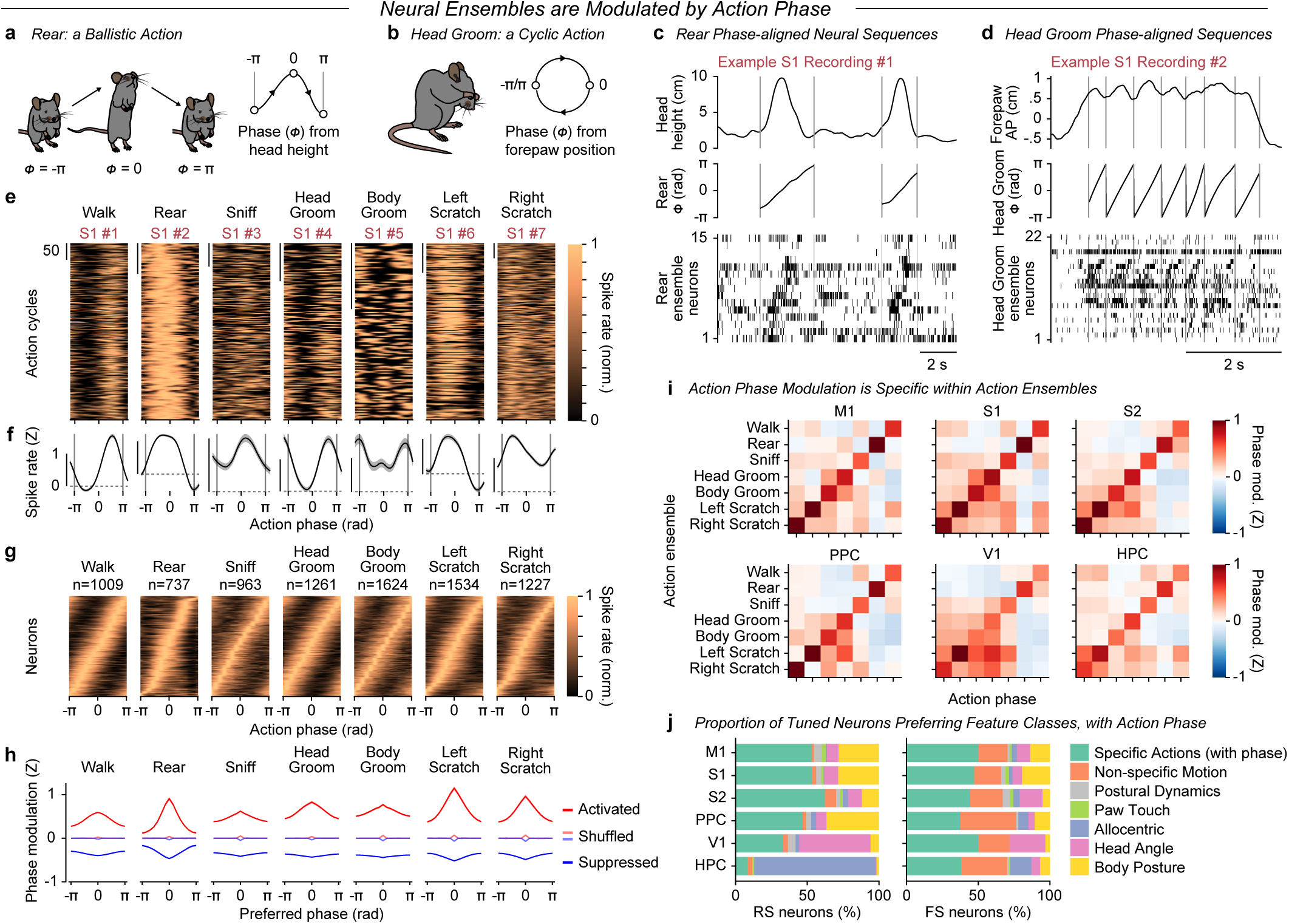
Action ensembles encode action phase as neural sequences. **a**, Illustration of the ballistic Rear action, with phase defined by head height. **b**, Illustration of Head Groom action, with phase cycle defined by forepaw anterior-posterior (AP) position. **c**, Example S1 recording containing rears. Top, head height; middle, Rear phase (*Φ*); bottom, raster with 15 neurons in Rear ensemble. Vertical gray lines mark Rear onset/offset. **d**, Same as (e), but for example S1 recording containing a head grooming bout. Top, mean forepaw AP trace; middle, Head Groom phase; bottom, raster from 22 neurons in the Head Groom ensemble. **e-f**, Peri-event time histograms (PETHs) aligned to action phase cycles for seven example S1 neurons tuned to each phasic action. PETHs are normalized within each cycle (e), and average PETHs (f) are standardized, with shading ±SEM. **g**, Action phase tuning curves for neurons in each activated action ensemble concatenated across cortical regions, normalized for each neuron and sorted by preferred phase. **h**, Average phase modulation (Z-scored), centered on each neuron’s preferred phase, for action-activated (red) and suppressed (blue) neurons, with shuffled controls for activated (light red) and suppressed (light blue) ensembles. **i**, Phase modulation grids for all combinations of action ensemble (rows) and action phase (columns) for each cortical region. **j**, Distribution of preferred feature classes for RS (left) and FS (right) neurons across cortical regions. Same format as Fig. 4h, but recalculated after including Action Phase in the Specific Actions feature class.

To determine whether these phasic action dynamics are reflected in cortical activity, we defined the relative position within each action cycle using Hilbert transforms to compute action phase (see Methods). This framework allows for phase alignment across repetitions of the same action, even if executed with varying speeds or durations (e.g., fast versus slow Rears, or different Walking gaits). When neural activity was aligned to action phase, striking temporal structure emerged: neurons within a given action ensemble consistently fired at the same phase across repetitions of the action cycle (e.g., Fig. 5c,d; see Supplementary Videos 2-6 for example ensembles tuned to different actions).

To validate this phase-specific firing pattern, we computed peri-event time histograms (PETHs) aligned to action phase onsets and offsets. Across repetitions, individual neurons showed highly consistent modulation at a specific phase within the action cycle (Fig. 5e,f). Remarkably, when phase tuning curves from all neurons in an action ensemble were pooled across brain regions, they formed a smooth sequence across the population that tiled the full action phase cycle (Fig. 5g; see region-specific heatmaps in Extended Data Fig. 9a). Similarly, neurons that were suppressed during actions exhibited phase-aligned suppression sequences, suggesting coordinated inhibition (see Extended Data Fig. 6g).

To further validate these phase tiling patterns, we quantified phase modulation by centering each neuron’s tuning profile on its preferred phase and computing the average Z-scored tuning across neurons within each action ensemble. This analysis, performed separately for activated and suppressed populations, revealed strong modulation around the preferred phase across each action ensemble (Fig. 5h; see region-specific phase modulation in Extended Data Fig. 9b). In contrast, phase modulation was negligible in shuffled controls. Moreover, phase modulation was action-specific: neurons in one action ensemble showed markedly reduced modulation by the phase of other actions across SMC, PPC, and even HPC (Fig. 5i).

Notably, the sequential firing patterns observed in action ensembles are unlikely to result solely from somatotopically-ordered cutaneous inputs. While tactile input may serve as a salient cue to anchor the neural sequence to certain phases of an action (e.g., forepaw contact at the onset or offset of a rear, brief whisker-forepaw contact during head grooming), touch events are unevenly distributed across the action cycle (see Extended Data Fig. 8c,d). If sequential patterns of tactile input were the primary driver of these sequences, we would expect tuning to certain phases associated with tactile events (Emanuel et al., 2021) to be overrepresented and other phases (i.e., lacking tactile input) to be underrepresented. In contrast, the phase tuning we observed smoothly tiled the entire action cycle (Fig. 5g).

Given the prominent action phase tuning observed in cortical neurons, we once again re-evaluated all recorded neurons to assess which feature class best explained their activity, now including action phase as a feature in the Specific Action class. The results revealed that for a stunningly large proportion of both RS and FS neurons, action phase emerged as the top explanatory feature (Fig. 5j), highlighting its role as a key organizing principle of cortical dynamics during natural behavior. To further assess how neurons jointly encode continuous and discrete aspects of actions, we decomposed the Specific Action category into three constituent variables: action phase, speed, and laterality (left-right direction). Across cortical areas, this revealed clear regional and cell-type specializations (Extended Data Fig. 9c,d). RS neurons in SMC were predominantly tuned to action phase, whereas FS neurons exhibited broader tuning to action phase and speed. PPC neurons showed a notable bias toward action laterality, consistent with their preference for midline turning postures. V1 and HPC populations, in contrast, showed weaker tuning to these action variables. Together, these analyses confirm that cortical populations encode combinations of continuous kinematic variables and discrete actions, providing a unified framework that links posture, motion, and action sequence representations. Thus, action ensembles serve as a neural substrate that supports not only the awareness of which action is being performed, but also awareness of the current action state, including phase, speed, and direction.

### A kinematic encoder model reveals neural subspaces for distinct actions

To test whether actions are indeed a dominant organizing principle underlying cortical dynamics, we developed an encoder model to predict population activity from continuous kinematic features, without providing any explicit information about action labels or phase. To this end, we implemented a modular EDTCN architecture incorporating 1) a shared “Core” kinematic encoder, trained across all sessions to learn a common set of latent features from input sequences of kinematic inputs (174 total features comprised of joint angles, head angles, body segment positions, and their temporal derivatives), and 2) a session-specific “Readout” layer that maps these shared features onto the activity of simultaneously recorded neurons (Fig. 6a; see Extended Data Fig. 10a). Our “Core+Readout” model outperformed other encoder architectures in predicting neural activity (Extended Data Fig. 10b,c; Methods), with especially strong performance in SMC and PPC. On held-out test data, the model explained substantial variance in RS neuron activity (Fig. 6b) across S1 (24.6%), S2 (15.7%), M1 (31.4%), and PPC (31.0%). In contrast, prediction performance was notably lower in V1 (10.3%) and HPC (13.1%), even though allocentric features including locomotion, place, and head direction (yaw) were among the input variables. Beyond predicting single-neuron activity, the model recapitulated the complex population dynamics observed in the recorded data (e.g., Fig. 6c).

**Figure 6.**
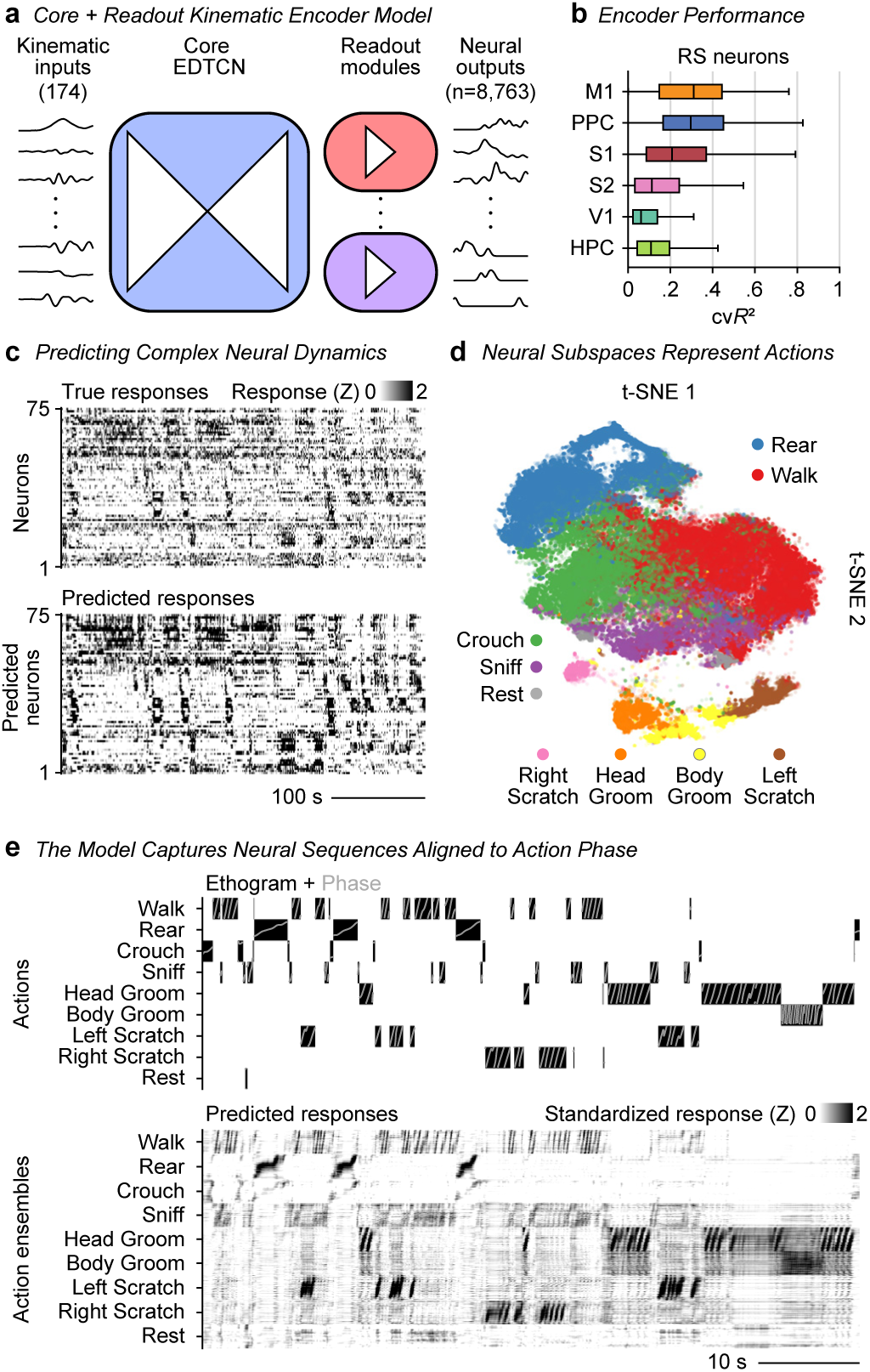
A kinematic encoder model recapitulates action sequence ensembles. **a**, Architecture of the Core+Readout kinematic encoder model. A batch of kinematic features, with bidirectional sequence (64 frames) is inputted to a shared encoder–decoder temporal convolutional network (EDTCN “Core”), producing 384 latent features. Latent features are then mapped through session-specific Readout layers to reduce the sequence dimension, each projecting into the neurons recorded in that session (total n=8,763 neurons across 392 sessions; number of neurons varies by session). During training, input features and Readout mappings were repeated for all sessions, allowing the Core to learn a shared kinematic representation, while each Readout module captured the neural translation. **b**, Box-and-whisker plot quantifying Core+Readout encoder model cross-validated fit (cvR²) for RS neurons recorded in each cortical area. Boxes represent interquartile ranges, lines mark median cvR², and whiskers span the full distribution. **c**, Example session showing true neural responses (top; Z-score normalized to 0-2 s.d.) and predicted responses (bottom; same normalization and sorting by Rastermap), capturing much of the dynamics in recorded data. **d**, t-SNE embedding of the predicted population activity (n=8,763 neurons). Dots represent time points, colored according to the nine actions classified by an independent action segmentation model. **e**, Example epoch showing the action ethogram overlaid with action phase (top) and predicted responses (bottom) for action ensembles (Walk: n=992; Rear: n=713; Sniff: n=949; Head Groom: n=1,207; Body Groom: n=1,561; Left Scratch: n=1,488; Right Scratch: n=1,220 neurons; Total, n=4,972 neurons). Neurons are sorted by preferred phase tuning (as in Fig. 5g). Predicted responses are Z-score normalized to 0-2 s.d. to emphasize activations.

Critically, the Core encoder applies 1-D temporal convolutions to sequences of kinematic inputs, such that predictions depend on a finite temporal context (effective receptive field of 640 ms) rather than a single frame. No action labels or action phase were provided to the model, ensuring that all predictions arose from continuous kinematic variables alone to uncover the organizing principles governing cortical activity. When we aggregated the model outputs into a pseudo-population of 8,763 neurons, sorting with Rastermap revealed distinct functional clusters that aligned to specific actions (Extended Data Fig. 10d). To visualize clustering at the population level, we applied t-SNE to the modeled neural activity from an example recording session (see Methods). Timepoints were colored by the corresponding action, revealing that the high-dimensional neural population vector segregated into discrete subspaces, each corresponding to a distinct action (Fig. 6d). Moreover, when the modeled neural activity was sorted into action ensembles based on empirical action and phase tuning, the modeled population exhibited clear sequential activity aligned to action phase (Fig. 6e). Together, these modeling results suggest that cortical dynamics during natural behavior organize population activity into action-specific subspaces and represent the state within the action cycle as sparse, phase-aligned ensemble sequences. These findings demonstrate that cortical neurons compress complex kinematic information into an efficient body schema centered around actions.

## Discussion

Here, we show that the cortex compresses high-dimensional kinematics in a compact, structured, and ethologically grounded representational format that underlies the body schema and supports action awareness. By combining full-body kinematic tracking with large-scale in vivo electrophysiology in freely moving mice, we discovered that the cortex encodes posture using distinct reference frames, integrates gravitational context, and organizes neural activity into action sequence ensembles. Together, these principles provide a foundation for how proprioceptive inputs are integrated into a coherent internal representation of the body in space, with implications for restoring sensorimotor function in cases of injury, stroke, or neurological disease.

Understanding the body schema requires identifying its representational format, particularly the reference frames in which it is encoded. Prior work (Wolpert & Ghahramani, 2000; Alonso et al., 2023) proposed egocentric reference frames for the posture representation. However, based on our findings during natural behavior, the body schema is anchored to gravity and the body’s midline (a hybrid egocentric-gravicentric coordinate system). We found that PPC primarily encodes postural angles relating to left-right turning (i.e., body midline reference frame), while SMC predominantly encode vertical height and pitch of the body with respect to the ground (i.e., gravicentric reference frame). This compact, physically-grounded coordinate system for representing 3D body posture allows the cortex to decompose high-dimensional kinematic inputs into ethologically-relevant components distributed across anatomically distinct regions. Furthermore, our systematic approach confirmed that V1 encodes posture in a head-centered reference frame (Mimica et al., 2023), and that the hippocampus encodes location and locomotion in an allocentric, world-centered reference frame (Moser et al., 2017).

To navigate and interact effectively with the physical world, land-dwelling vertebrates must anchor their posture representation relative to the ground. Crucially, “ground” is not fixed—it can be the flat surface of an arena, a tree branch, or the floor of a tunnel—so the brain must maintain an internal reference rooted at the body’s base of support and aligned with gravity to stabilize posture across diverse gaits and terrains. The “idiotropic vector”, a concept introduced by Mittelstaedt, refers to this centrally stored, gravity-aligned axis that anchors the perception of the upright direction and supports spatial orientation across contexts (Mittelstaedt, 1999). Our findings provide evidence in support of this conceptual model, suggesting that SMC integrates tactile input from ground contact with proprioceptive and vestibular cues to construct body-part positions in a gravicentric coordinate system. This representation enables rapid readout of body elevation relative to the ground, supporting efficient postural control. Unlike joint-centric angles, gravity-referenced features, such as elevation and pitch, provide a global representation of posture within the gravitational field. Our findings support the view that sensorimotor integration in SMC contribute to a grounded, gravity-referenced body schema. This reference frame enables the brain to monitor and adjust posture to support balance, facilitating stable motor control across diverse environments and behavioral demands. Future work is needed to determine how sensorimotor and parietal areas integrate multisensory cues to construct and maintain this gravity-and ground-referenced body schema.

From an evolutionary perspective, it is adaptive to represent midline-referenced body configurations separately from ground-referenced postures. The observed overrepresentation of PPC neurons tuned to tail and trunk angles, particularly those associated with left-right deviations from the midline, raises the possibility that PPC is a body-centric state estimator that predicts the postural and sensory consequences of turning, with a special role in compensating lateral balance during axial rotations. During turning movements, animals must generate preemptive adjustments to maintain balance, such as initiating compensatory tail movements before the onset of whole-body turns. In this context, encoding of tail abduction-adduction and roll angles in PPC may reflect predictive motor representations that help coordinate such adjustments. Supporting this idea, PPC neurons in rodents encode upcoming turns during maze navigation, consistent with a broader role in predictive coding of future actions (Harvey et al., 2012; Whitlock et al., 2012). Our findings that the PPC encodes posture and actions relative to the midline is consistent with clinical and experimental observations of hemi-spatial neglect, in which PPC lesions impair awareness of the contralateral side of the body (Brain, 1941; King & Corwin, 1993; Karnath et al., 2002). Thus, PPC appears to contain an internal forward model that predicts the physical consequences of turning actions, enabling smooth, balanced, and coordinated voluntary movements.

During natural behavior, the brain must not only track posture but also anticipate the body’s biomechanics across different action phases. The varying gravitational demands across phases likely contribute to the formation of distinct action ensembles. For example, during an unsupported rear, the rear-up phase requires muscular effort to raise the body’s center-of-mass against gravity. The hold phase demands postural stability under sustained gravitational load, as the animal balances upright with its full weight supported by the hindlimbs. In the rear-down phase, gravity assists the descent, but engagement of forelimb and trunk reflexes upon ground contact must absorb impact and restore balance in a quadrupedal stance. More broadly, each distinct action places unique biomechanical demands across different phases, imposing a need for phase- and action-specific representations and downstream recruitment of spinal and brainstem circuits (Wang et al., 2017; Yang et al., 2023). Across all cortical areas examined, we found a substantial fraction of neurons exhibited highly stable, phase-locked firing patterns aligned to action cycles. Hence, rather than encoding static postures, these neurons preferentially represented transitions between dynamic posture changes, aligned to distinct action phases.

Our findings also suggest that the cortex compresses the high-dimensional space of dynamic postural configurations into more compact representations that reflect the stereotypical posture and movement patterns associated with individual actions. Supporting this idea, our modeling showed that population activity is organized into low-dimensional subspaces associated with distinct actions. Thus, the brain effectively addresses the challenge of representing the vast space of kinematic configurations in a freely moving, multi-jointed body. Natural behavior thus constrains this kinematic space based on natural statistics, which the brain efficiently encodes as action subspaces. These action subspaces parallel continuous attractor models in which continuous variables, such as head orientation or spatial location, are encoded as localized activity clusters on a low-dimensional manifold embedded within a high-dimensional neural state space (Kakaria & de Bivort, 2017; Khona & Fiete, 2022; Kutschireiter et al., 2023). While previous studies reported sequential neural dynamics during isolated, constrained behaviors such as treadmill locomotion (Foster et al., 2014; Xing et al., 2022; Borgognon et al., 2025) or a trained reaching task (Churchland et al., 2012; Grier et al., 2025), our findings unveiled the prevalence of distinct action ensembles whose sequential activity tiled the full phase cycle of multiple species-typical actions during freely moving behavior. Our observation that each action recruits distinct neural sequences reveals a previously underappreciated organizational principle of cortical dynamics during natural behavior (Miller et al., 2022).

Our behavioral paradigm captured a broad range of spontaneous full-body actions, but it did not encompass the full richness of the mouse behavioral repertoire. Previous studies utilizing a fine forelimb behavior such as skilled reaching, manual manipulation, or climbing revealed cortical representations with activity aligned to reach phase (Wang et al., 2017; Yang et al., 2023; Grier et al., 2025) or reaching direction (Galiñanes et al., 2023). Future work in enriched environments and social or task-motivated contexts may reveal additional action ensembles or finer-grained structure within action subspaces (Marshall et al., 2021; Klibaite et al., 2025). Nevertheless, our framework predicts that any species-typical action would recruit distinct cortical ensembles. Phase-aligned activation patterns within action ensembles reflected the stereotyped structure of natural behavior, where both posture and neural activity unfold in predictable sequences. Specific neural sequences may encode upcoming postural transitions, rather than current or past sensory states, thus enabling a predictive internal model of motor intention and upcoming sensory consequences. Along the same lines, sequential dynamics could also simplify transitions between actions, such as rearing to walking, providing a circuit-level substrate for the planning of voluntary actions to support smooth continuity across behaviors.

Low-intensity microstimulation of human PPC elicits an urge to move without overt movement, consistent with biasing latent action circuits rather than directly driving muscles (Desmurget et al., 2009). In line with these observations, we found action-aligned dynamics in PPC. Ensemble recruitment could underlie the conscious experience of actions, and cortical microstimulation in PPC may recruit these action ensembles to produce the illusion of action. A subset of interneurons in PPC, as well as SMC, showed action-specific, phase-aligned responses. Inhibitory interneurons may contribute to network-level gating, suppressing competing ensembles and allowing only the selected action sequences to unfold. Furthermore, many putative interneurons showed strong correlations with global body motion. Recurrent activity among excitatory and inhibitory neurons in these network ensembles could update the body schema at the proper rate as actions are performed at various speeds (e.g., fast versus slow rearing or walking). This evidence suggests motion-related excitation and inhibition, along with neuromodulatory tone (Pakan et al., 2016), could synchronize and set the pace of action-sequence progression across the cortex.

Together, our findings uncover core organizing principles underlying the body schema during spontaneous behavior. By revealing how neural ensembles in the cortex encode posture and action, we provide a foundation for understanding the neural basis of the body schema and action awareness. These insights could inform future brain-machine interfaces and bio-inspired robotics, where embedding ground- and gravity-referenced sensing may improve body-part localization, postural control, and autonomous movement planning. Finally, integrating physically realistic biomechanics and physics-based simulations (Aldarondo et al., 2024), together with action context into dynamical systems models may enable more adaptive and flexible motor control across diverse tasks and environments.

## Methods

### Animal care

Adult C57BL/6J (JAX stock #000664) mice were obtained from the Jackson Laboratory. During electrophysiological and behavioral experiments, mice were single housed under a 12-hour normal light/dark cycle with ad libitum access to food and water. All experiments were conducted during the light phase. Details regarding the sex and age of all animals are provided in Supplementary Table 1. All procedures were performed in accordance with protocols approved by the Institutional Animal Care and Use Committees at the Massachusetts Institute of Technology.

### Procedures

Mice were implanted with head-posts to fix the head during optical imaging, microdrive implantation, and headstage connection. Mice were anesthetized with isoflurane (3 % for induction, 1.0-1.5 % for maintenance) delivered in oxygen at 0.8 L/min and positioned in a stereotaxic frame (Model 963, David Kopf Instruments). A custom stainless steel headbar (Pololu; 0.060″ ± 0.004″, Grade 304) was affixed to the skull using cyanoacrylate adhesive (Krazy Glue, #KG585) and sealed with a protective layer of dental cement (Stoelting).

Targeting of visual cortex (V1) was performed stereotaxically to coordinates 3.5 mm posterior, and 2.5 mm lateral to Bregma on the right hemisphere. For all other target areas, targeting was aided by intrinsic optical signal (IOS) imaging to localize hemodynamic responses to vibrotactile stimulation of different body parts. Following full recovery from headpost implantation, mice were lightly anesthetized with isoflurane (0.5–1%). Kwik-Cast and dental cement covering the skull were removed to provide optical access. Body temperature was maintained at 37°C with a heating pad (FHC), and respiration was continuously monitored. Illumination was delivered using high-power LEDs (Thorlabs M530L3, green for vascular reference; M625L4, red for hemodynamics) oriented at oblique angles onto the cortical surface. Reflected light was collected by a tandem lens macroscope (objective lens: Canon 50mm f1.2; imaging lens: Nikon 85mm f1.4) and imaged by sCMOS camera (PCO Edge 4.2). Images were acquired at 20 Hz at 512×512-pixel resolution (with 4x average binning) using PCO acquisition software. Vibrotactile stimulation (Seeed, Part #316040001; 4s, 10s inter-stimulus interval) was applied to the skin and/or fur of the contralateral neck, back, tail, forepaw, hindpaw, or E-row whiskers. White noise auditory stimulation was presented continuously to minimize auditory cortex activation. Reflectance changes (ΔR/R) were calculated by normalizing pixel intensities to pre-stimulus baselines and averaging across trials (≥10 repetitions). Post hoc analysis aligned regions-of-interest showing stimulus-locked decreases in reflectance with vascular landmarks to guide subsequent craniotomies for tetrode microdrive implantation.

Prior to microdrive implantation surgery, mice were anesthetized with isoflurane (3 % for induction, 1.0-1.5 % for maintenance) delivered in oxygen at 0.8 L/min and secured in a stereotaxic frame (Model 963, David Kopf Instruments). Preoperative analgesia (meloxicam, 0.5 mg/kg, subcutaneous) was administered to reduce inflammation. A ground screw was inserted into the contralateral hemisphere and pinned to the EIB. Following a small craniotomy and durotomy at the target area, the tetrode bundle was lowered just below the brain surface. Moveable components of the microdrive and the craniotomy were coated with sterile ophthalmic ointment (Puralube). The microdrive was affixed to the skull with dental cement. A shield, made of transparency film laminated with copper tape and connected to the EIB, was cemented to the skull and covered with black electrical tape to protect the implant during postoperative recovery. Postoperative analgesia (buprenorphine, 0.05 mg/kg, subcutaneous) was administered on the day of surgery. Animals were monitored and received continued analgesia (meloxicam, 0.5 mg/kg, subcutaneous) for at least two additional days and were allowed to recover for a minimum of seven days prior to recording.

### Electrophysiology

For microdrive construction, tetrodes were prepared by clamping four insulated nichrome alloy microwires (Sandvik, #PX000003) into a tetrode spinner (NeuraLynx, version 2.0). In manual mode, the wires were rotated 100 turns clockwise, followed by 40 counter-clockwise turns to release residual torsion. The tetrode bundle was heat-treated with a heat gun (350–400 °C, low setting) held 2 cm from the wires for 5 seconds from three directions to fuse the bond coat while preserving the insulation. Individual tetrode wires were pinned by gold pins (Neuralynx, #31-0603-0102) to a custom 64-channel electronic interface board (EIB, designed in DipTrace and printed by OSH Park) equipped with two 32-channel Omnetics connectors. The fused end of each tetrode was routed through a polyimide tube (ID: 99 μm, OD: 168 μm, Polymicro Technologies) arranged in a 2×8 or 4×4 array and epoxied to the nano-Drive (Cambridge Neurotech). Once routed, tetrodes were glued to the polyimide array and trimmed to equal length using tungsten carbide scissors (FST, #14502-14). Prior to implantation, tetrodes were immersed in gold-plating solution (Neuralynx, #31-0611-0100) and electroplated using Nano-Z 1.2 (White Matter) to a target impedance of 200 kΩ. In some cases, tetrodes were coated with the fluorescent dye DiI (Invitrogen, #V22885) to assist in post hoc histological confirmation of recording sites.

For chronic electrophysiological recordings, mice were briefly head-fixed to minimize mechanical strain while connecting the 2×32 channel Omnetics connectors on the 64-channel headstage (Intan, Part #C3325). The brain was allowed to relax for at least 15 minutes before transferring the animal to the denoised recording arena, enclosed within a grounded Faraday cage. The headstage was tethered via a 6-ft ultra-thin SPI cable (Intan, Part #C3216) to a custom passive commutator, allowing unrestricted movement during recording sessions. Extracellular voltage signals were acquired at 30 kHz using an RHD2000 Acquisition Board (Intan, Part #C3100) and RHX Data Acquisition Software (Intan). Signals were hardware filtered between 0.1 Hz to 10 kHz to enable separation of fast- and regular-spiking neuronal populations based on waveform shape. At the end of each session, mice were briefly head-fixed to allow safe removal of the headstage and advancement of the tetrode array by 62.5 µm increments. Recording sites were verified by post hoc histology.

For spike sorting, initial automated single-unit isolation was performed using Kilosort 3.0 (https://github.com/MouseLand/Kilosort). Tetrodes were mapped onto 16 “shanks” of four channels each. Extracellular voltage signals were high-pass filtered at 300 Hz and whitened prior to automated clustering in Kilosort3 using rigid drift correction (nblocks = 1). Clusters were subsequently manually curated in MClust 4.4 (http://redishlab.neuroscience.umn.edu/MClust). Isolation quality was assessed using: false alarm (FA) rate, defined as the fraction of spikes with inter-spike intervals (ISI) shorter than 2 ms (< 2% threshold); mean firing rate (FR > 0.25 sp/s threshold); and signal-to-noise ratio (SNR > 5 s.d. threshold, defined as mean waveform amplitude divided by the standard deviation of baseline noise for the peak channel). Units not meeting these criteria were excluded from further analysis. Units with a narrow waveform shape (trough-peak duration < 0.45 ms) were labeled as putative fast-spiking (FS), and units with a wide waveform shape (trough-peak duration > 0.45 ms) were labeled as putative regular-spiking (RS).

### Histology

Mice were deeply anesthetized and transcardially perfused with phosphate-buffered saline (PBS; Gibco), followed by 4% paraformaldehyde (PFA) in PBS. Brains were post-fixed overnight and then cryoprotected in 30% sucrose in PBS at 4°C. Tissue was embedded in OCT compound (Sakura Finetek) and coronally sectioned at 80 μm using a cryostat (Leica Biosystems). Sections were stained with Neurotrace Blue (1:500; Thermo Fisher Scientific, #N21479) in 0.3% Triton X-100/PBS. The sections were then briefly rinsed with PBS and mounted on slides with a custom-made mounting medium (Takatoh & Prevosto et al., 2021). All histological images were acquired with a Keyence microscope (BZ-X800). Locations of recording tracks were manually registered to the Allen CCF.

### Behavioral recordings

The recording arena was constructed on a 1 × 1 m anodized aluminum breadboard (Thorlabs, Part #MB30) mounted on rubber feet (Thorlabs, Part #RDF1) to minimize vibration. The enclosure was assembled using a custom T-slotted aluminum frame with grounded aluminum side panels to provide electrical shielding. The arena floor consisted of a multi-layered base (foam underlay, PVC baseboard, and green vinyl film) designed to provide traction and durability. A cylindrical glass enclosure (hurricane candle holder; inner diameter 189 cm, height 205 cm) surrounded the arena, allowing unobstructed visual access. Illumination was provided by three to four grounded 30 W LED arrays (IKAN), ensuring uniform lighting conditions. Between recording sessions, both the glass cylinder and arena floor were cleaned by wiping with 70% ethanol.

Six machine-vision color cameras (acA1920-150uc; Basler) were operated via Pylon Software (Basler) to record video at 1152 × 1024 resolution in RGB pixel format using on-chip PGI 5 × 5 debayering. Cameras acquired frames with a global shutter synchronized by external hardware frame triggers at 100 fps with 2 ms exposure time. Each camera was mounted with an 8 mm fixed-focal-length lens (M0824-MPW2, Computar) with 300 mm working distance. Polarizing filters (PR120-30.5; Midwest Optical) were mounted on the lenses to reduce glare and reflections from the glass cylinder and other reflective surfaces. Cameras were mounted using ½″ posts and accessories (Thorlabs) and arranged radially around the arena at ∼60° intervals. To provide sufficient parallax for 3D triangulation, cameras were angled vertically at alternating inclinations of 0° (horizontal) and −15° (downward).

Intrinsic camera calibration (lens distortion and focal parameters) was performed using a 10 × 11 cm checkerboard. Extrinsic calibration (camera pose and orientation) was conducted using a custom 3D-printed L-frame affixed to a circular plexiglass base (8″ diameter, 0.25″ height). Camera parameters were computed using custom MATLAB scripts and achieved sub-pixel reprojection error for each view.

Multi-camera triggering and video recording were managed by Campy python software (https://github.com/ksseverson57/campy). Camera triggering was synchronized by a microcontroller (Teensy 3.2) running custom Arduino triggering software (Campy trigger module). Video frames were acquired using custom Python recording software (CamPy), with each stream compressed in real time using FFmpeg configured with H.264 or H.265 NVENC hardware encoding on NVIDIA GPUs. Multi-camera video files were acquired as single MP4s. Total video acquired were reduced in size from ∼10 TB/hr to ∼10 GB/hr when compressed at 4 Mb/s constant bit rate, making saving, storing, and loading video datasets manageable while preserving quality (visually lossless). To enable precise alignment of video frames with electrophysiological recordings, trigger TTL pulses were simultaneously recorded on analog input channels of the RHD2000 board at 30 kHz sampling rate.

### Markerless 3D pose estimation

DANNCE pose estimation models were architected and trained similarly to procedures described in Dunn, Marshall et al., 2021. Briefly, a total of 835 frames from 15 sessions were manually annotated using the Label3D GUI to define 3D positions for a 44-keypoint mouse skeleton. 775 frames were used for training, with 60 frames held out for validation. A 2D convolutional neural network (CNN) was first trained to estimate the center of mass (COM), defined as the geometric mean of the labeled keypoint positions. The 3D CNN DANNCE model was finetuned from pre-trained weights originally trained on the Rat7M dataset (Dunn & Marshall et al., 2021). During DANNCE model finetuning, data augmentation included random jittering of the centroid position, rotation, and mirroring (with left-right keypoint re-indexing) of the 3D volumetric grid to improve generalization. Synchronized frames from the six cameras were projected into the augmented 80×80×80 volumetric grid (1.5 mm isometric voxel resolution) using projective geometry. Additional augmentations included random variation in view selection, brightness, and hue. To improve smoothness of keypoint predictions, a static “weighted average” layer on top of the “MAX” network outputted the weighted average position of the probability map. Keypoint prediction accuracy was evaluated on a held-out validation set (n=60 frames) by computing the 3D Euclidean distance between predicted and ground truth keypoint positions. Once validated, the trained COM and DANNCE models were applied to all behavioral recordings (340* sessions across 18 mice, totaling 122.4 million frames). Keypoint positions were smoothed using a 5-frame or 11-frame moving average filter for segment/joint calculations and tuning analyses, respectively.

### Geometric full-body kinematics

We developed a geometric anatomical framework to quantify full-body kinematics with high precision. This geometric anatomical model transforms 3D keypoint tracking data (x,y,z) into angles defined in world-, body-, and joint-centered reference frames (Extended Data Fig. 1). A total of 26 anatomical keypoints were tracked across the body to capture the position and rotation of each major segment and joint, with at least three keypoints per segment to resolve 3D plane orientation. Joint-centered axes were aligned to the anatomical planes of each segment: the anteroposterior axis (X) followed the proximal-distal orientation, the mediolateral axis (Y) aligned to segment width, and the vertical axis (Z) was defined as the cross-product of X and Y, capturing long-axis rotation. Using this geometric body model, we defined rotations for 16 major skeletal joints (i.e., neck, thoracic-lumbar and lumbar-sacral spine, tail, shoulders, elbows, wrists, hips, knees, and ankles) encompassing 38 rotational degrees of freedom. This comprehensive framework provides a continuous, anatomically-defined description of the body configuration in freely moving animals, enabling detailed parameterization of body posture, motion, and postural dynamics.

For allocentric angle calculation, rotation matrices for each body segment were constructed from the X, Y, and Z axis vectors derived from 3D Cartesian keypoint positions. Allocentric rotation angles (yaw, pitch, roll) were then computed relative to the calibrated world coordinate frame using a “YXZ” Euler angle sequence. This sequence was selected over the more common “XYZ” convention to avoid gimbal locks, as pitch angles in our mouse behavior recordings occasionally exceed ±90° (e.g., during high rearing or grooming), while head roll rarely exceeds 90°. Subsequent transformations, including body-centered alignments, were computed using an “XYZ” Euler sequence applied to the allocentric angles.

For joint-centered angle calculation, joint rotation matrices were computed by multiplying the rotation matrix of the distal (target) segment by the transposed rotation matrix of the proximal (basis) segment. This “rotation-of-coordinates” convention preserves the hierarchical structure of the skeleton, allowing angles to propagate from proximal to distal joints and enable full reconstruction of 3D keypoint positions via forward kinematics. The resulting joint rotation matrix (*Rj*) captures the relative rotation of the distal segment with respect to the proximal segment at joint *j*. To maximize the dynamic range of each joint and avoid gimbal locks, we centered the rotation at the mean or “T-pose” orientation for each joint by multiplying the inverse of the singular value decomposition (SVD) of the average joint rotation matrix with the joint rotation matrix. With this SVD T-pose alignment procedure, Euler sequence choice was less impactful.

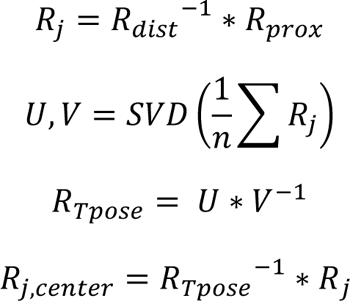

Joint-centered Euler angles were then extracted from these rotation matrices using axis sequences selected to yield interpretable components: flexion–extension (*fe*), abduction–adduction (*aa*), and internal–external rotation (*ie*). For each joint, we validated the selected Euler sequence by binning angles and visualizing average aligned keypoint positions across bins to confirm correspondence to expected biomechanical axes (Extended Data Fig. 1). To account for biomechanical constraints, special considerations were made: elbow and knee joints were modeled as single-axis hinge joints and restricted to *fe* only, computed via trigonometric angle between vectors; tail joint angles were restricted to *fe* and *aa*, as *ie* rotation could not be estimated with a single midline keypoint on the tail. In total, 38 joint angles were calculated. Angles were smoothed using either a mild moving average (21-frame sliding window) or Savitzky-Golay filter (fifth order, 21-frame sliding window) applied to unwrapped angles to reduce noise while preserving high-frequency postural dynamics. Parallelized computation using MATLAB’s “pagemtimes” function enabled efficient, frame-independent Euler angle extraction. The precision of joint angle error calculations was estimated via Monte Carlo simulation by injecting random Gaussian noise (from keypoint Euclidean error in Extended Data Fig. 1b) into predicted keypoint positions (n = 3,600 frames) and re-computing segment and joint rotations (n = 1,000 repetitions).

We performed principal component analysis (PCA) on a combined set of joint angles and head and body segment orientation angles to identify low-dimensional representations of full-body posture. Angles were first centered in circular coordinates, and outliers were removed using percentile-based filtering. Wrist joints were excluded due to nosier tracking. Features from 37 full-length (one hour) recording sessions were concatenated and Z-scored prior to PCA. PCA was performed on the pooled feature matrix to identify dominant orthogonal postural modes (PCs), and cumulative variance explained was used to assess the dimensionality of the posture space. PC loadings were saved and projected onto additional sessions; for example, posture PC scores were used to assess neural tuning by correlating firing rates with percentile-binned PCs. To visualize posture dimensions captured by individual PCs, timepoints were binned into quintiles along each PC axis, and the mean 3D keypoint positions were computed for the lowest and highest quintiles. Pose data were smoothed and aligned to a common reference frame (lumbar spine segment). The mean pose was rendered in 3D using a predefined keypoint skeleton. This procedure revealed interpretable low-dimensional posture motifs along PC axes, such as vertical elevation (Z), forward and backward leaning (X), and leftward or rightward turning (Y). Body motion was quantified as the smoothed mean angular speed averaged across all joints, excluding wrists and ankles to reduce noise. Angular velocities were clipped to the 1st–99th percentile range to mitigate outliers, smoothed using a 21-frame moving average, and expressed in degrees per second. Locomotion speed was defined as the magnitude of frame-to-frame head displacement in the horizontal (XY) plane, also smoothed with a 21-frame moving average and expressed in millimeters per second.

To estimate proximity (touch) of body parts to either the ground or other body parts as proxies for touch, we computed two features: distance-to-floor (d2f) and distance-to-face (d2face). D2f features were computed as the minimum Euclidean distance between each paw segment and the floor or wall, whichever was closer, clipped at 5 mm. D2face features were computed as the Euclidean distance between each paw and the snout keypoint, as well as the distance between each paw and the ipsilateral ear, clipped at 5 mm. Derivatives of d2f and d2face were computed to estimate touch dynamics.

For raster plotting, Rastermap was used to sort neuron order (Stringer et al., 2025). Briefly, spike rates were computed from spike counts using a 40-ms Gaussian convolution. Spike rates were Z-scored and inputted into Rastermap (https://github.com/MouseLand/rastermap) for single neuron sorting with the following settings: n_PCs = n_neurons/2, locality = 0.5, time_lag_window = 100 frames (1 second), symmetric = False. For rasters showing activations, spike rates were Z-scored, sorted by the Rastermap neuron order, plotted as an image, and color limited to positive range to emphasize activations.

### Tuning curve calculation

To quantify stimulus selectivity, we computed marginal (1D) tuning curves by estimating each neuron’s conditional firing rate as a function of each continuous behavioral or kinematic variable. Firing rates for tuning curve calculation were computed by convolving 1-ms binned spike counts with a 40-ms Gaussian kernel. Stimulus variables were first filtered to exclude outliers using percentile-based thresholds. For most features, stimuli were binned into 25 equal-percentile bins. Categorical variables (e.g., action category, action phase) were handled separately using discrete levels, while uniformly spaced binning was applied to touch-related variables (e.g., distance to floor, distance to face), head direction, and place (in 5 × 5 grid). For each bin, the average spike rate and SEM were computed across timepoints assigned to that bin. Tuning curves were generated only for bins with at least 50 samples. To assess tuning reliability, we computed cross-validated R² values using a five-fold data split following the partitioning strategy described above. Mutual information (MI) between stimulus and response was calculated from the bin-wise firing rate conditional expectations and the stimulus distribution. As a control, shuffled tuning curves were generated by randomly permuting spike-stimulus pairings, and 95% bootstrap confidence intervals were computed across resampled datasets.

To perform feature selection using marginal tuning models to identify the most informative kinematic- or action-related feature for each neuron, we applied marginal tuning models across an extensive feature set (294 total features). For each neuron-feature pair, we computed the tuning curve and assessed its predictive power using cross-validated R². Stimuli were binned before computing the conditional mean firing rate in each bin, which served as a lookup table to predict responses on held-out data. Predictions were then compared to actual firing rates, and R² was calculated as the proportion of variance explained (see Encoding Models below). This process was repeated across all candidate features, and the feature yielding the highest cross-validated R² was selected as the neuron’s best-tuned feature. To avoid noisy estimates from low sample counts, negative or out-of-bound R² values were clipped between 0 and 1. Neurons with no marginal tuning model *R*² above 0.05 (5% variance explained, approximately 0.5 bits/s, see Extended Data Fig. 3) were considered “untuned”. This method provided a simple, robust, and interpretable estimate of top feature importance for each neuron in the recorded population.

Kinematic features were grouped into functional categories to aid in interpretability of tuning analyses. Posture features included static joint angles, body segment pitch and roll angles, and segment heights; Postural dynamics included first and second derivatives of posture features (head, body segment, and joint angular velocity and acceleration, lateral and vertical motion and acceleration); Head angles included head pitch and head roll angles; Touch-related features included proximity measures (e.g., distance-to-floor, distance-to-face); Allocentric (world-centered) features included place and head direction; Movement features included whole-body motion metrics (mean joint speed, locomotion speed). Action features included coarse action categories classified by the supervised action segmentation model (see below). The full list of features and their assigned classes is provided in Extended Data Fig. 3f.

### Action segmentation

Ground truth action annotations were manually classified by human labelers using a modified version of the DeepEthogram GUI (Bohnslav et al., 2021), adapted to display multiple camera views simultaneously. Short video epochs were selected to represent each major action class observed in our mouse behavior recordings (see definitions below). Each frame was assigned a single, mutually exclusive label corresponding to one of eight primary action categories: walk, rear, crouch, sniff, head groom, body groom, scratch, or rest. Frames containing infrequent behaviors, such as wet dog shake, forepaw shake, or stretch, were labeled as other. A total of 477,372 frames (79.6 minutes) were annotated from six mice (3 male, 3 female). Final labels were exported from the GUI as a one-hot logical matrix in CSV format, enabling framewise accuracy calculation for classifier evaluation on held-out test partitions (see below).

We used the following operational definitions of species-typical actions: *Walk*: Alternating forepaw and hindpaw movements for forward locomotion and/or turning. *Rear*: Both forepaws lifted off the ground and the torso height >40% of maximum. *Crouch*: Intermediate torso height (<40%) without other active behaviors. *Sniff*: Either (a) prone posture (resting) with snout lifted to sniff air, or (b) prone posture (walking) with head tilted downward and snout paused close to the ground to sniff floor or consume food. *Head Groom*: Use of forepaws to clean the head (ears, face, whisker pad, snout, mouth) or use of the tongue to lick the glabrous side of the forepaw between head grooming bouts. *Body Groom*: Crouched posture with head directed to groom the body (arms, abdomen, back, legs) with mouth, or grooming hindpaws using forepaws and mouth; also includes licking the dorsal surface of the forepaw. Scratch: Hindpaw lifted to rapidly claw (∼10-20 Hz) at the head or body, or brought to the mouth to lick between scratching bouts. *Rest*: Sedentary periods in prone posture with low movement speed. *Other*: Rare behaviors (e.g., wet dog shake, forepaw shake, stretch) comprising ∼1% of annotated actions.

To perform unsupervised action segmentation from 3D keypoint data, we implemented keypoint-MoSeq (Weinreb et al., 2024) with custom adaptations. Because the original pipeline was designed for DeepLabCut keypoints, we developed a data loader to import DANNCE-generated 3D keypoint coordinates, converting them into a compatible format. We specified anterior and posterior body parts for egocentric alignment and excluded tail keypoints to focus on core body and limb movements. The model was trained on six hours of data (2,160,000 frames) from six mice (2 male, 4 female). PCA was used for initialization, retaining components that captured 90% of variance. To achieve a median syllable duration of ∼300 ms, we tuned the stickiness parameter (κ) over a range from 1×10^3^ to 1×10^15^, settling on κ = 9×10^9^. After 50 AR-HMM initialization steps, the full model was fit using 500 iterations of Gibbs sampling, with convergence tracked via log joint probability. The final model identified 95 unique syllables (e.g., “Syllable 0” … “Syllable 94”). After visual inspection of short behavior video snippets, we assigned semantic labels to each syllable and grouped similar syllables into 25 intermediate clusters (e.g., Rear Up, Rear Down), which were then mapped to nine coarse behavior categories (Walk, Rear, Crouch, Sniff, Head Groom, Body Groom, Left Scratch, Right Scratch, and Rest). For syllables with overlapping features (e.g., Rear and Sniff), we assigned the category with higher priority. Model accuracy was evaluated on a separate set of human-annotated frames excluded from training. The unsupervised clustering provided by keypoint-MoSeq was instrumental in discovering the coarse action categories and correlations between neural activity and actions.

We refined the action segmentation labels with a supervised model. We implemented a custom AS-EDTCN model in PyTorch, adapted from a previously described action segmentation architecture (Lea et al., 2017). This encoder-decoder network compresses temporal kinematic dynamics into a low-dimensional representation via an encoder block, and the decoder block outputs categorical action labels. The encoder consists of two blocks of 1D convolutional layer followed by layer normalization, ReLU activation, max pooling, and dropout for regularization. The decoder mirrors this structure, with upsampling, dropout, 1D convolution, layer normalization, and ReLU activation. The model processes an acausal bidirectional input sequence of 128 frames, downsampled 2x, resulting in a total temporal receptive field of ± 1280 ms. The temporal resolution was downsampled via max pooling across two encoder blocks and restored the original sequence length through two decoder blocks via upsampling. A static attention mechanism, implemented as a trainable einsum operation, reduced the sequence dimension to a single output frame. The predicted label was assigned as the class with the maximum value (argmax). Categorical cross-entropy loss with class weighting was used to prevent overfitting to more frequent action classes. Full training hyperparameters included: AdamW optimizer, exponential learning rate decay with warm-up (γ = 0.87), batch size = 64, pool size = 4, sequence length = 128, hidden sizes = [384, 384], kernel size = 8, learning rate = 1×10^-6^, L2 regularization = 1×10^-6^, elastic net ratio (L1:L2) = 0.01:0.99, and dropout = 0.25. Generalization performance was evaluated using leave-one-subject-out validation, ensuring that test data from the unseen subject were held out from training.

Accuracy of the supervised (AS-EDTCN) and unsupervised (Keypoint-MoSeq) action segmentation models were assessed by averaging the per-class true positive rate (TPR) across all action classes in held-out test data. Specifically, the supervised AS models were trained on labeled data from five out of six subjects and tested on the remaining subject to assess generalization. Unsupervised action labels predicted by the Keypoint-MoSeq model were validated on the same labeling dataset (N = 6 subjects). The true positive rate (TPR), was defined for each class as the proportion of correctly classified positive examples relative to all actual positive examples:

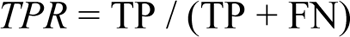

Here, TP (True Positive) refers to instances where the action was correctly predicted, FN (False Negative) refers to instances where the action occurred but was incorrectly classified as another action. In parallel, the false positive rate (FPR) was computed to quantify the proportion of negative cases (i.e., timepoints not belonging to the given action class) that were incorrectly classified as positive, where FP (False Positive) refers to timepoints wrongly predicted as belonging to the class, and TN (True Negative) refers to timepoints correctly predicted as not belonging to the class:

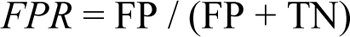

Overall classification accuracy was calculated as the average TPR across all classes:

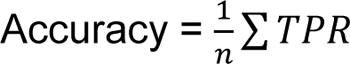

To account for the non-uniform distribution of action classes, the cross-entropy loss during supervised training was weighted by the inverse class frequency. This class-balancing approach prevents over-weighting common actions and under-weighting rare actions, thereby ensuring that accuracy metrics reflect performance across the full range of actions.

### Power spectral density analysis

To identify frequency-domain signatures of specific behaviors, we computed the power spectral density (PSD) of kinematic features across stereotyped actions. Kinematics from a subset (N=21) recording sessions were aggregated, including keypoint positions and action label identified by our AS-EDTCN classifier (e.g., grooming, scratching, rearing). Kinematic signals were first preprocessed by subtracting a local moving average (midpoint filter) to emphasize relatively fast movements. The resulting signals were then bandpass filtered (0.1–20 Hz) using a third-order Butterworth filter. For each behavior and feature (e.g., head height, forepaw height, etc.), PSDs on 1-dimensional features were estimated using Welch’s method (256-point window, 128-point overlap, 512-point FFT) with a Hamming window. PSDs were averaged across behavior episodes and plotted as frequency-by-feature matrices to visualize unique kinematic “fingerprints” associated with each action class.

### Action phase calculation

To estimate the continuous phase (i.e., polar position within the action cycle) of ongoing actions, we manually selected a representative kinematic feature for each action whose dynamics reflected its stereotyped, periodic trajectory. These features were chosen based on prior PSD analysis and neural correlations to capture the dominant motion and neural encoding associated with each action. Each selected feature was bandpass filtered using a third-order zero-phase Butterworth filter, with filter parameters tuned to the dominant frequency range of each action. The instantaneous phase was computed using the Hilbert transform, yielding continuous action phase ranging from −π to π, defined only within bouts of the corresponding action:

Walk: difference in hindpaw anteroposterior (AP) position;

1-3 Hz Rear/Crouch: head height; 0.1-2 Hz

Sniff: head pitch; 2-8 Hz

Head groom: mean forepaw AP position;

1-3 Hz Body groom: inverted head height;

2-8 Hz Left/Right scratch: left/right hindpaw height;

1-3 Hz Rest: not assigned (not phasic)

To discretize these continuous phase signals, the phase cycle of each action was divided into three equal-width phase bins corresponding to the rising, peak, and falling phases of the stereotyped trajectory. This procedure produced 3×N+1 total bins for the N = 8 phasic actions, with an additional bin assigned to “rest”, resulting in 25 discrete action phase bins.

To identify functional ensembles of neurons tuned to specific actions, we classified neurons based on their action-specific modulation strength using Z-scored tuning values. Tuning to coarse action categories (N=9 categorical bins) was first computed (see *Tuning Curve Calculation*), and the motion-independent component of tuning (motion-subtracted residual action tuning) was used for thresholding. We computed action modulation as the total modulation of the Z-scored action tuning curve (max-min). This approach allowed us to categorize neurons into ensembles for each action. A neuron was classified as “activated” by a particular action if its positive Z-scored modulation was greater than a positive threshold (+0.25 s.d.), “suppressed**”** if its negative modulation was below a negative threshold (−0.25 s.d.), or “modulated” if the total modulation exceeded the threshold (0.25 s.d.). We opted for an effect size measure because the statistical power of our large sample size (360,000 frames) makes even the weakest correlations statistically significant. This effect size threshold method has the advantage of non-exclusive ensemble membership, and thus a neuron could be part of multiple ensembles if it exhibited strong modulation to more than one action. This contrasts with clustering methods, which typically enforce mutually exclusive assignment and may obscure shared functional networks across actions.

### Action phase tuning analysis

To analyze phase-aligned peri-event time histograms (PETHs), we aimed to characterize the temporal relationship between neural activity and ongoing cyclic actions. We computed phase-aligned PETHs by aligning neural activity and stimulus features to the onset and offset of actions. We then aligned spike rates and characteristic action features (e.g., head height, hindpaw height) to individual phase cycles of the action, enabling averaging across repetitions despite variable action durations. Continuous spike rates were estimated from binned spike counts using a Gaussian smoothing kernel with width equal to half the phase cycle duration. Spike rates were normalized by z-scoring the log-transformed firing rates (using the numerically stable log1p function) across the session. Action cycles were first segmented based on the action category and further separated into distinct action cycles based on action phase (see Action Phase Calculation). For each cycle, the representative stimulus feature (e.g., hindpaw AP position for walking, head height for rearing) was extracted and time-warped by interpolating the phase cycle duration to the target duration (100 samples). The corresponding spike rate trace was also interpolated across the same normalized phase axis using linear interpolation. Trials were thus aligned and binned into a fixed number of phase points per cycle (e.g., 100 phase bins ranging from −π to π) to enable direct averaging across cycles. Phase-aligned PETHs were computed separately for each action class and neuron, allowing fine-grained dissection of firing patterns across stereotyped action cycles.

To demonstrate tiling of action phase tuning, we sorted neuron order within each ensemble (e.g., activated, suppressed, modulated) according to their preferred phase. Phase tuning curves were extracted for each neuron using the Z-scored firing rate across discrete action phase bins (25 total; see Action Phase Calculation), separately for each action. Tuning curves were upsampled by a factor of 10 and smoothed to reduce sampling noise and allow finer sorting resolution. Neurons were sorted based on their peak or trough response phase, depending on ensemble type (e.g., peak for activated, trough for suppressed). Interpolated tuning curves were Z-scored per neuron, normalized to the [0,1] range, and plotted as heatmaps sorted by preferred phase. Neuron IDs for each ensemble, sorted by preferred phase, were used to sort predicted population activity (see below).

To quantify action phase modulation, we computed the Z-scored tuning curve for each neuron and extracted the amplitude of modulation across the action cycle. For each action and neuron, Z-scored tuning curves were aligned to the preferred phase using a circular shift. Total phase modulation was quantified as the difference between the maximum and minimum response across the phase cycle. Modulation was also computed for shuffled data, in which spike rates were computed from randomly permuted spike counts, providing a baseline for comparison, as alignment to peak phase will always produce a sequence, albeit with minimal modulation. Finally, for each ensemble and region, we computed an action–phase modulation matrix capturing how, on average, neurons in each action ensemble modulated activity across all other action phase cycles to evaluate phase-specific ensemble recruitment.

### Kinematic encoder models

We performed static data partitioning to avoid temporal auto-correlation. Each one-hour recording session was divided into ten continuous 6-minute segments (“deciles”). For five-fold cross-validation, each decile was split into 80% training and 20% test data. To prevent temporal overlap, a three-second buffer around each training–test boundary was excluded from both sets, following the approach in Syeda et al., 2024. Partition masks were fixed (non-random) and reused consistently across all encoding model training and evaluation procedures.

Regression model (neural encoder) performance was assessed by computing the coefficient of determination (*R*^2^) on held-out test data, where *y* is the true response, *ŷ* is the predicted response, and *ӯ* is the mean response:

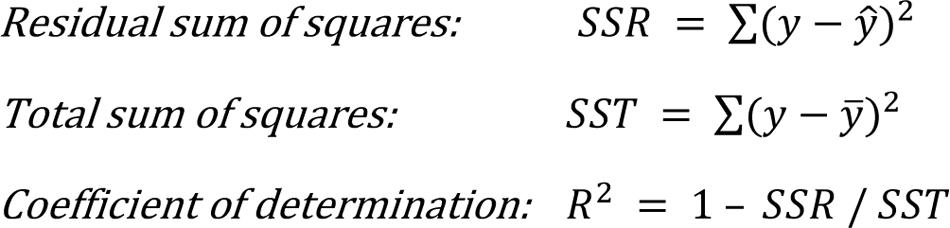

We trained a core–readout encoder-decoder temporal convolutional network (EDTCN) to predict neural activity solely from kinematic features (with no action information). In benchmark testing, this architecture outperformed alternative models, including GLMs, multilayer perceptrons (MLPs), gradient boosted decision trees (XGBoost), Kolmogorov-Arnold networks (KANs), RNNs, LSTMs, and various CNN architectures on a neural regression (encoder) task. The EDTCN model consisted of a shared temporal convolutional encoder (“Core”) that transformed sequences of kinematic inputs into a lower-dimensional latent representation, and a set of session-specific fully connected decoders (“Readouts”) that mapped this latent activity onto the responses of individual neurons. The encoder was comprised of an input bidirectional 1D convolution (kernel size = 8, dropout = 0.25, hidden size = 384), and two additional convolutional layers, each followed by layer normalization, ReLU activation, and max pooling (pool size = 2). The decoder block consisted of two feedforward layers of upsample (scale factor = 2), dropout, 1D convolution, and ReLU (see schematic in Extended Data Fig. 10). Kinematic features were inputted into the shared EDTCN “Core” as sequences with a total temporal receptive field of ±640 ms, and included joint angles, head orientation, head/trunk/paw positions, and their temporal derivatives (velocity and acceleration), as well as place, locomotion, and body motion (total 174 kinematic features). Action labels were excluded. These kinematic features were Z-scored per session using statistics from the training set. Neural firing rates were normalized between 0 and 1. The model was trained across 340 recording sessions from multiple brain regions (M1, PPC, S1, S2, V1, HPC) using mini-batch gradient descent (AdamW optimizer; batch size=128; initial learning rate = 1 × 10⁻⁵; linear warmup schedule). Training was performed in mixed precision (FP16) and regularized by elastic net (L1:L2 ratio = 0.01:0.99). Model loss was computed as the root mean squared log error (RMSLE) between predicted and observed firing rates. To prevent overfitting, we used five-fold cross-validation with early stopping. Preprocessed data were stored as memory-mapped arrays to optimize memory usage. Final model accuracy was assessed using the coefficient of determination (*R*²), computed independently for each neuron.

### Predicted population analysis

We used a t-SNE embedding to perform dimensionality reduction on predicted population activity. To predict population neural activity from the kinematic inputs in an example session, we first applied the trained Core EDTCN module to predict latent representation, followed by a Readout layer concatenated across all recorded sessions, outputting a total of 8,763 RS neurons. The resulting population activity matrix (timepoints × neurons) was Z-scored across time and subjected to principal component analysis (PCA) for dimensionality reduction. The first ∼140 principal components (PCs), explaining at least 90% of the variance, were retained. These components were then embedded into two dimensions using t-SNE (https://github.com/pavlin-policar/openTSNE), with perplexity = 500 and default early exaggeration settings optimized for visualization. Each point in the resulting 2D embedding represents a population activity vector at a single timepoint. To visualize clustering with respect to action category, each point was color-coded by the action label for that frame, based on the automated AS-EDTCN classifier prediction. This procedure enabled visual inspection of action-specific structure in the predicted neural state space, capturing separable trajectories and clusters corresponding to species-typical actions.

### Statistics

To test whether EDTCN outperformed alternative encoder architectures, we performed paired, one-sided Wilcoxon signed-rank tests on per-neuron predictive performance (R²), comparing EDTCN > GLM and EDTCN > LSTM separately for RS and FS neurons. Neurons contributed one value per model after averaging 5-fold cvR² within neuron. Analyses were pooled across all cortical areas. Because R² distributions were skewed, we used nonparametric tests and reported effect sizes as the median ΔR² (EDTCN − comparator) with bootstrap 95% CI (2,000 resamples). Multiple comparisons were controlled using Benjamini–Hochberg False Discovery Rate (BH FDR) across the four pooled tests (RS/FS × GLM/LSTM); significance threshold *p* < 0.05. Extremely small p-values were truncated and reported as less than 1×10⁻¹⁶.

## Acknowledgements

We thank members of the Wang and Dunn labs, Antoine De Comite, Mehrdad Jazayeri, and Steve Flavell for valuable discussions and feedback. We thank Paul M. Thompson and Seonmi Choi for technical assistance. This work was supported by NIH grants DP1 NS137188 to F.W. and F32MH122995 to K.S.S.

## Author contributions

F.W. and K.S.S. conceived the project. K.S.S. and F.W. designed experiments with input from T.W.D. K.S.S. performed all experiments with assistance from W.X., H.J., and M.S.L. K.S.S. analyzed the data with help from J.L., H.J., T. Lou, M.S.L., and W.X. T.W.D., T. Li, and K.S.S. developed the DANNCE models. J.L. and K.S.S. developed the geometric anatomical framework. H.J., T. Lou, K.A.C., and W.X. assisted K.S.S. with action segmentation models. H.J., T. Lou, and T. Li assisted K.S.S. with the neural encoder models. K.S.S. and F.W. wrote the manuscript with input from all authors. F.W. supervised the project.

## Data availability

Source data is provided with the manuscript. Data is available in NWB (Neurodata Without Borders) format on the DANDI repository: dandiarchive.org/dandiset/001608 (access will be unembargoed upon publication). The pose estimation demo is available at github.com/ksseverson57/DANNCEv2. All other datasets, including raw multi-camera behavioral video data, will be made available upon reasonable request from the corresponding authors. Source data are provided with this paper.

## Code availability

All code used to reproduce figures in this paper and key analysis pipelines, including phase extraction, action segmentation, and neural encoding models, are available at github.com/wanglab-neuro/Severson_et_al_2025. The CamPy multi-camera acquisition software is available at github.com/ksseverson57/campy. All custom code is released under an academic, non-commercial research license that permits free use for research purposes but prohibits redistribution and commercial use. Full license terms are provided on the respective GitHub repository pages.

## Materials availability

All unique materials generated in this study (e.g., custom electronic interface board, headcap CAD files) are available from the corresponding author upon reasonable request. Commercial reagents and off-the-shelf components are listed with vendor, catalog numbers, and RRIDs (if available) in the Methods/Key Resources Table.

## Supplementary information

**Extended Data Figure 1.**
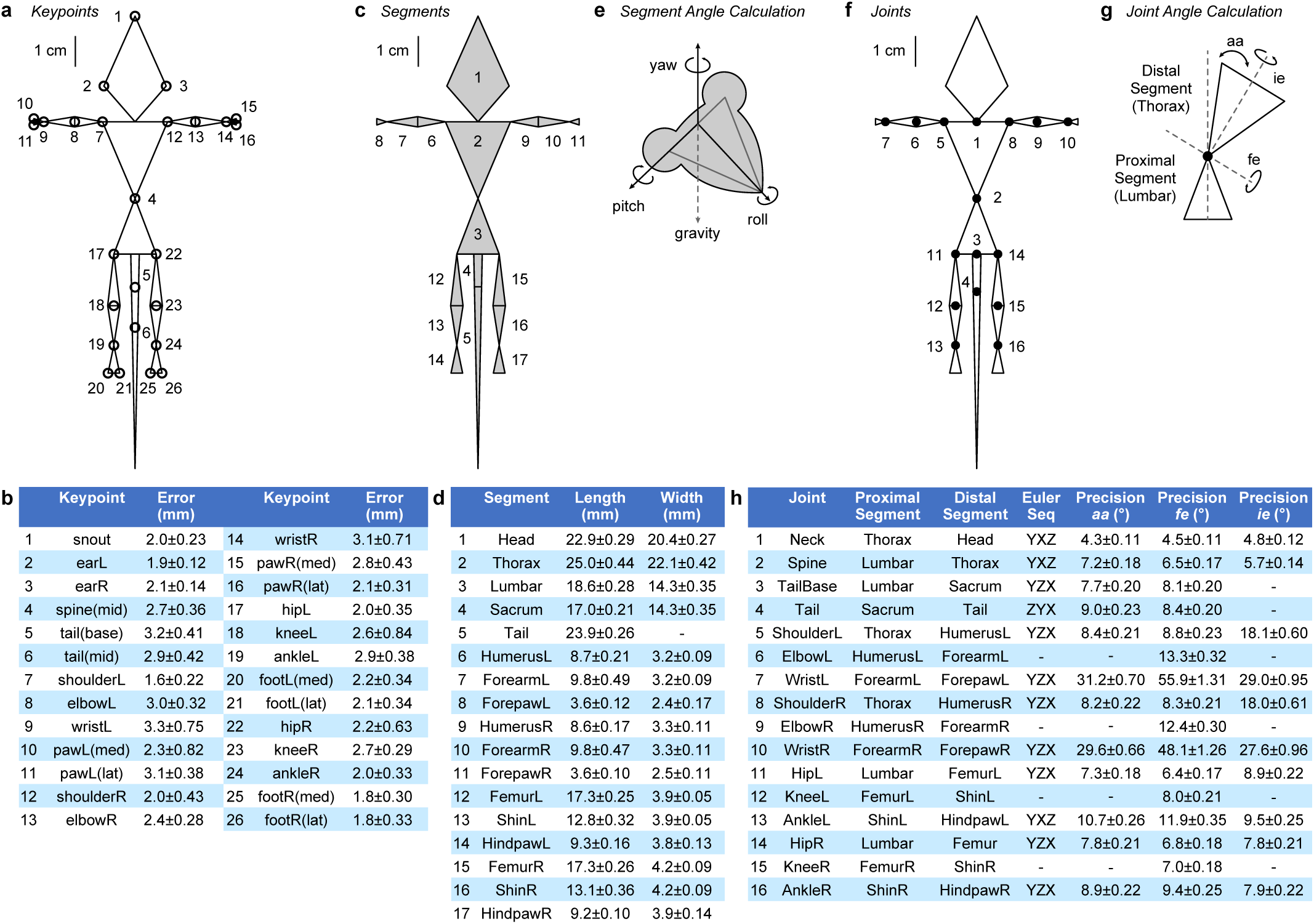
Anatomical keypoint, segment, and joint definitions for computation of segment and joint rotations. **a,** Scale rendering of 26 mean predicted anatomical keypoints (open circles) from an example mouse recording. **b**, Table defining the 26 anatomical keypoints used for computing body segment and joint angles. Error represents the mean Euclidean distance (±95% CI, in mm) between human annotations and DANNCE predictions on a validation dataset (60 frames from 12 labeled sessions). **c**, Scale rendering of 17 mean predicted body segments (gray polygons) from the same example recording in (a). **d**, Table defin-ing each the 17 body segments in the geometric model. Segment length and maximum width (±95% CI) were averaged across 17 mice. **e**, Schematic illustrating how segment angles were calculated relative to the global reference frame defined during extrinsic calibration, such that pitch and roll were measured with respect to gravity. **f**, Scale rendering of the 16 mean predicted joint positions (filled circles) and correspond-ing body segments (open triangles) from the same recording in (a) and (c). The 17 body segments act as rigid links connecting the 16 articulated joints comprising the anatomical skeletal model. **g**, Schematic illus-trating joint angle calculation using a three degree-of-freedom example joint (Spine). Flexion-extension (fe) angle measures the pitch, abduction-adduction (aa) angle measures the yaw, and internal external (ie) long-axis rotation angle measures the roll of the distal segment relative to the proximal segment. Euler angles were computed from the relative rotation matrices (Rdist and Rprox) using the Rotation-of-Coordi-nates convention and joint-specific Euler rotation sequence listed in (h). **h**, Table defining the 16 major joints, their proximal segment (toward body center), distal segment (toward extremities), Euler angle sequence used to compute the joint angles, and average precision error of joint angles associated with each joint, ±95% CI, in degrees. Joint angle precision was estimated via Monte Carlo simulation using keypoint error in (b).

**Extended Data Figure 2.**
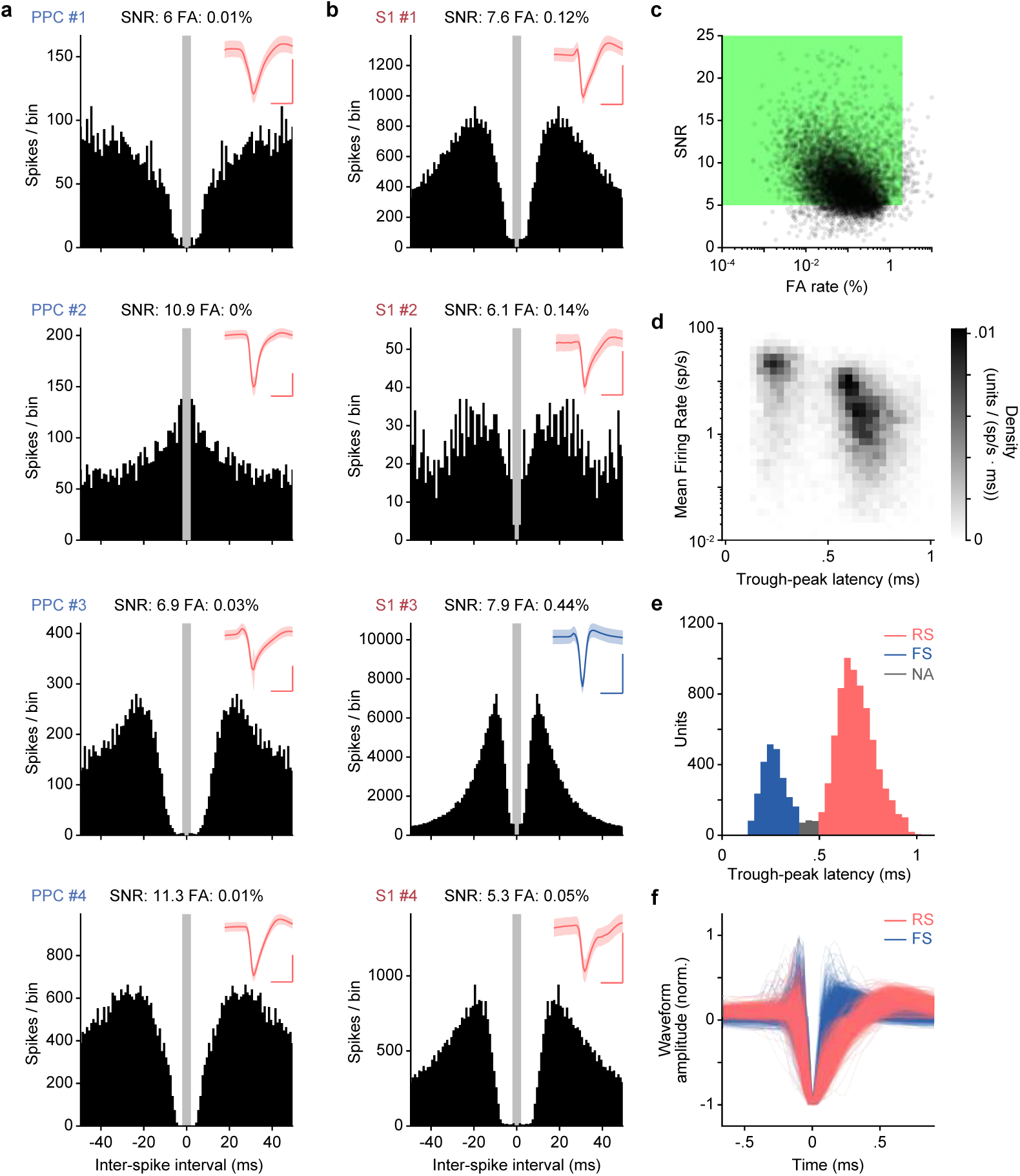
Spike sorting quality metrics. **a-b**, Inter-spike interval (ISI) histograms for example units from PPC (a, Fig. 2) and S1 (b, Fig. 3). Gray shaded region indicates ±2 ms refractory period. Insets show mean spike waveforms (RS, red; FS, blue); shaded areas represent ± s.d. Scale bars: 0.5 ms (horizontal), 0.2 mV (vertical). **c**, Scatter plot of wave-form signal-to-noise ratio (SNR) versus ISI false alarm (FA) rate for the recorded dataset (n=11,047 total units). The area within the green square contains acceptable units with SNR > 5 and false alarm rate < 2% (n=9,632 units from 18 mice). **d**, Density plot of units binned by mean firing rate and trough-peak latency, showing distinct clusters of putative interneurons (left; higher mean firing rate, narrow waveforms) and putative excitatory neurons (right; lower mean firing rate, wider waveform). **e**, Histogram showing bimodal distribution of trough-peak latencies used to separate FS (blue, latency < 0.4 ms) and RS (red, latency > 0.5 ms) neurons. Neurons with ambiguous waveform (NA, gray, latency 0.4-0.5 ms) were removed from subsequent analyses. **f**, Example typical waveform shapes (n=500) for units classified as FS (blue) and RS (red), normalized to trough amplitude.

**Extended Data Figure 3.**
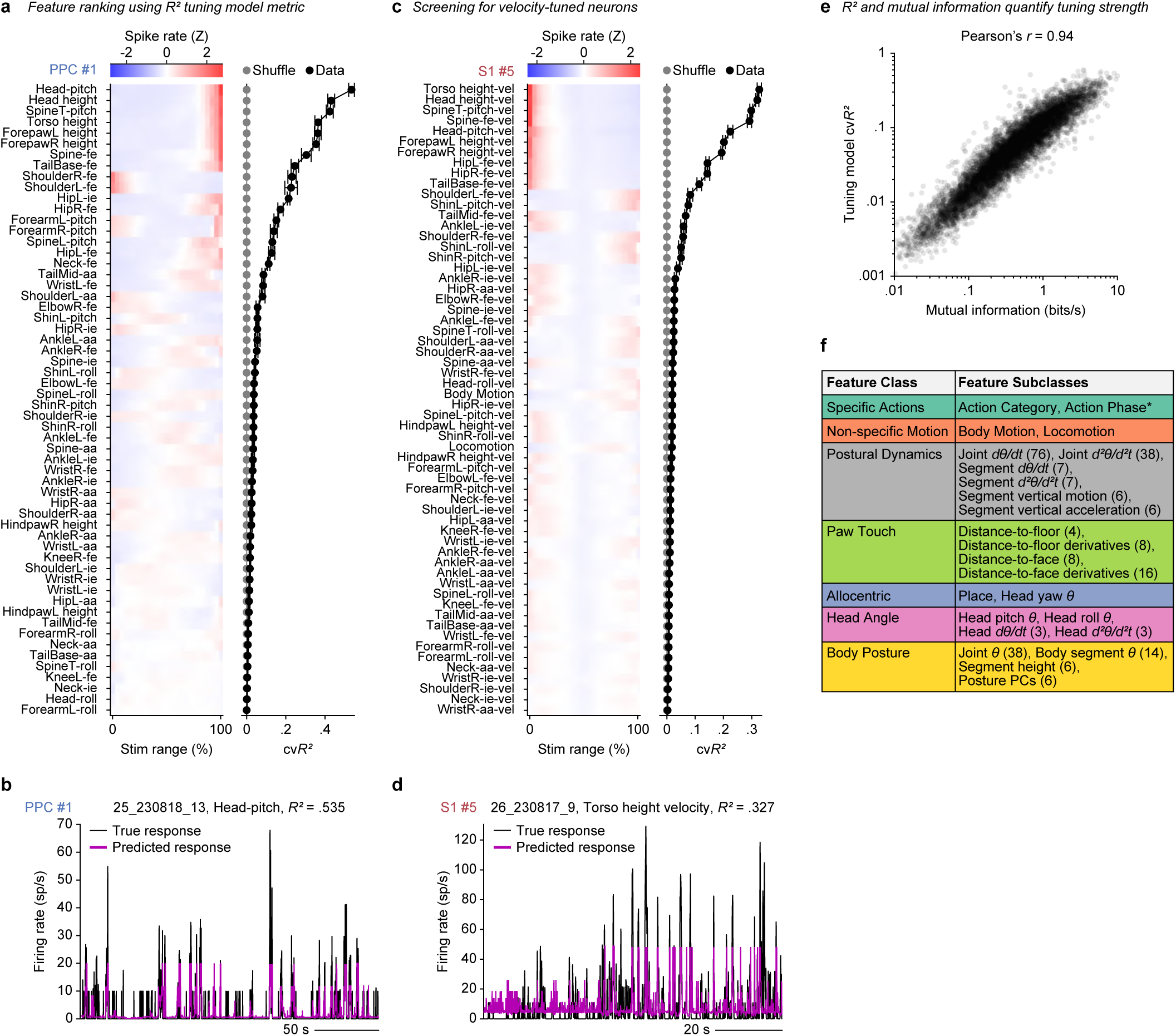
Feature selection using tuning models. **a**, Left, Z-scored tuning curves for example PPC neuron #1 across all posture features, sorted by ascend-ing cross-validated prediction performance (cvR²) of marginal (1D) tuning models. Right, cvR² for aligned data (black) and temporally shuffled data (gray; 30-min circular shift), shown in the same sorting order. Error bars represent ±SEM from 5-fold cross-validation. **b**, Predicted spike rate (magenta) versus observed firing rate (black) for the PPC neuron in (a). **c**, Same as (a), but for an example S1 neuron (#5, not shown in Fig. 3) tuned to posture dynamic features (e.g., change in torso segment height). **d**, Predicted (magenta) versus observed firing rate (black) for the S1 neuron in (c). **e**, Scatter plot comparing mutual information (MI) and cvR2 for each neuron’s best feature (n = 8,582 neurons), showing a strong correlation (Pearson’s r = 0.94, p < 1 × 10-7). **f**, Table of feature subclasses grouped into broader feature classes used to categorize neurons in Fig. 3t, Fig. 4h, and Fig. 5j. Numbers in parentheses indicate the number of features in each subclass. *, Action Phase used exclusively in Fig. 5j.

**Extended Data Figure 4.**
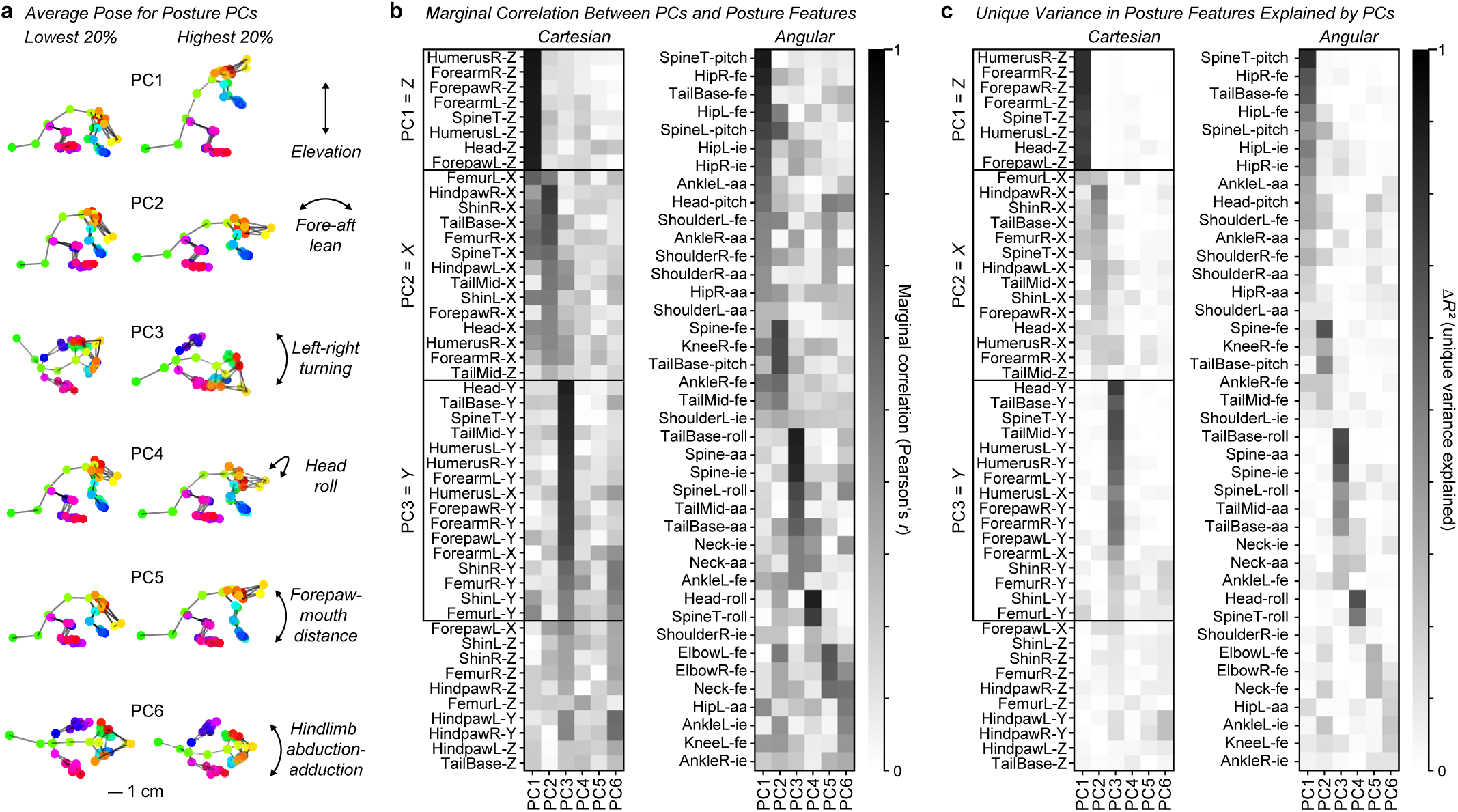
Eigenposes represent interpretable low-dimensional body posture modes. **a**, Average poses for the lowest and highest 20th-percentile bins from each of the first six Posture PCs from an example recording (different from Fig. 2n). **b**, Heatmap showing Pearson’s correlation between first six Posture PCs and two sets of features: left, Cartesian coordinates (XYZ positions); right, angular posture features. Features are sorted by the PC with the strongest correlation. **c**, Heatmap showing incremental variance in each Cartesian (left) or angular (right) feature explained uniquely by each Posture PC after controlling for other PCs.

**Extended Data Figure 5.**
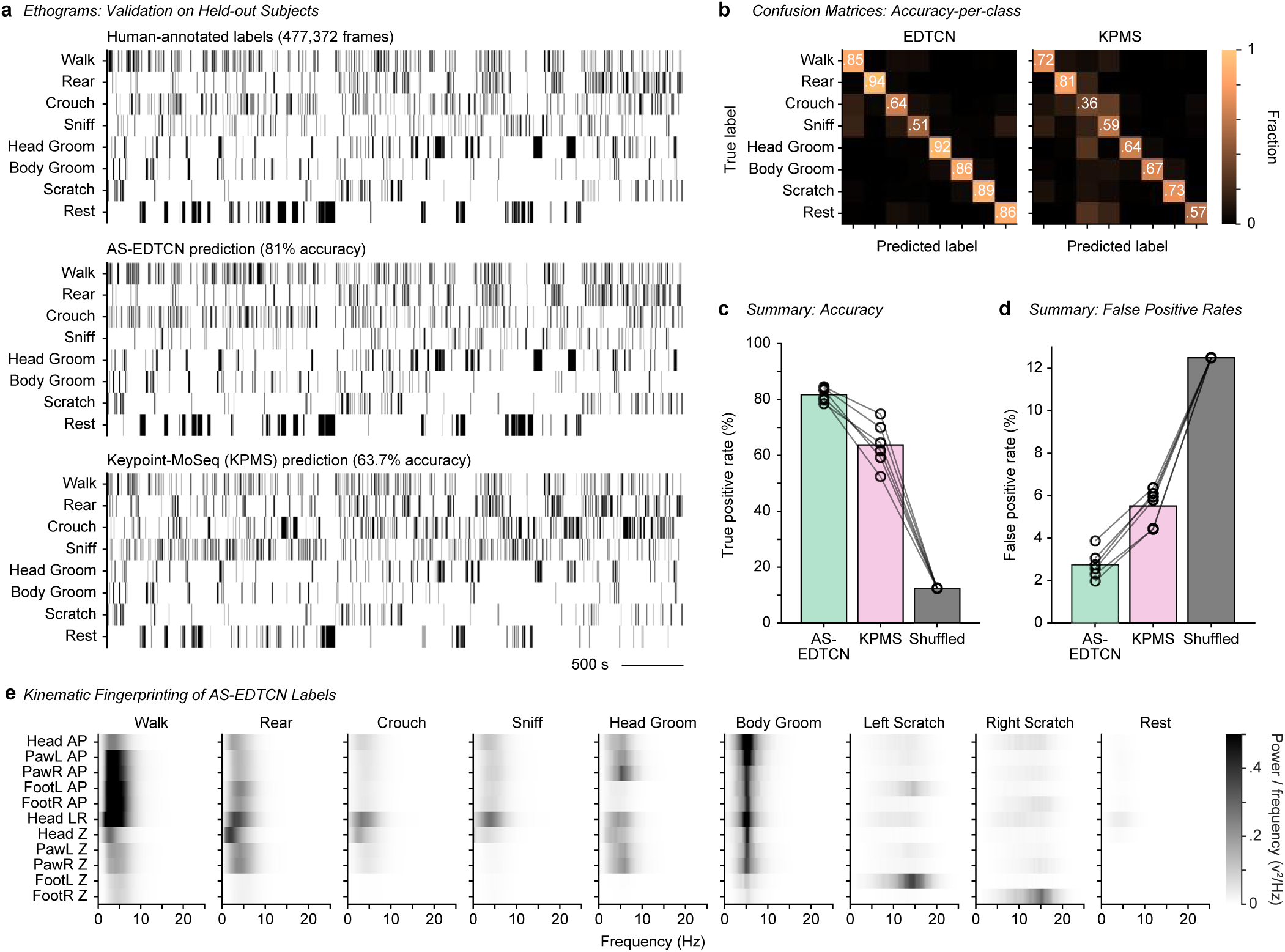
Validation of action segmentation models. **a**, Top, Ethogram of validation frames labeled by human curators, concatenated across six recordings from six mice (477,273 frames total). Middle, Predictions from the supervised AS-EDTCN model on held-out test data using a leave-one-subject-out cross-validation strategy (accuracy: 81.0%). Bottom, Predictions from the unsupervised Keypoint-MoSeq (KPMS) model on the same held-out test data (accu-racy: 63.7%). **b**, Left, Confusion matrix showing prediction proportions across true class labels for the AS-EDTCN model. Right, Average confusion matrix for the KPMS model on the same validation data. **c**, Class-balanced true positive rate (diagonal of the confusion matrix) for AS-EDTCN (green), KPMS (pink), and shuffled control (gray). True positive rates for individual recordings (N=6 mice) are shown as open circles connected by lines. **d**, Mean false positive rate for each model (chance=1/8 classes), overlaid with false positive rates for each action class as open circles (8 action classes). **e**, Power spectral density of body part position signals (y-axis) across frequency bands (x-axis), computed for samples in each action class as segmented by the AS-EDTCN model.

**Extended Data Figure 6.**
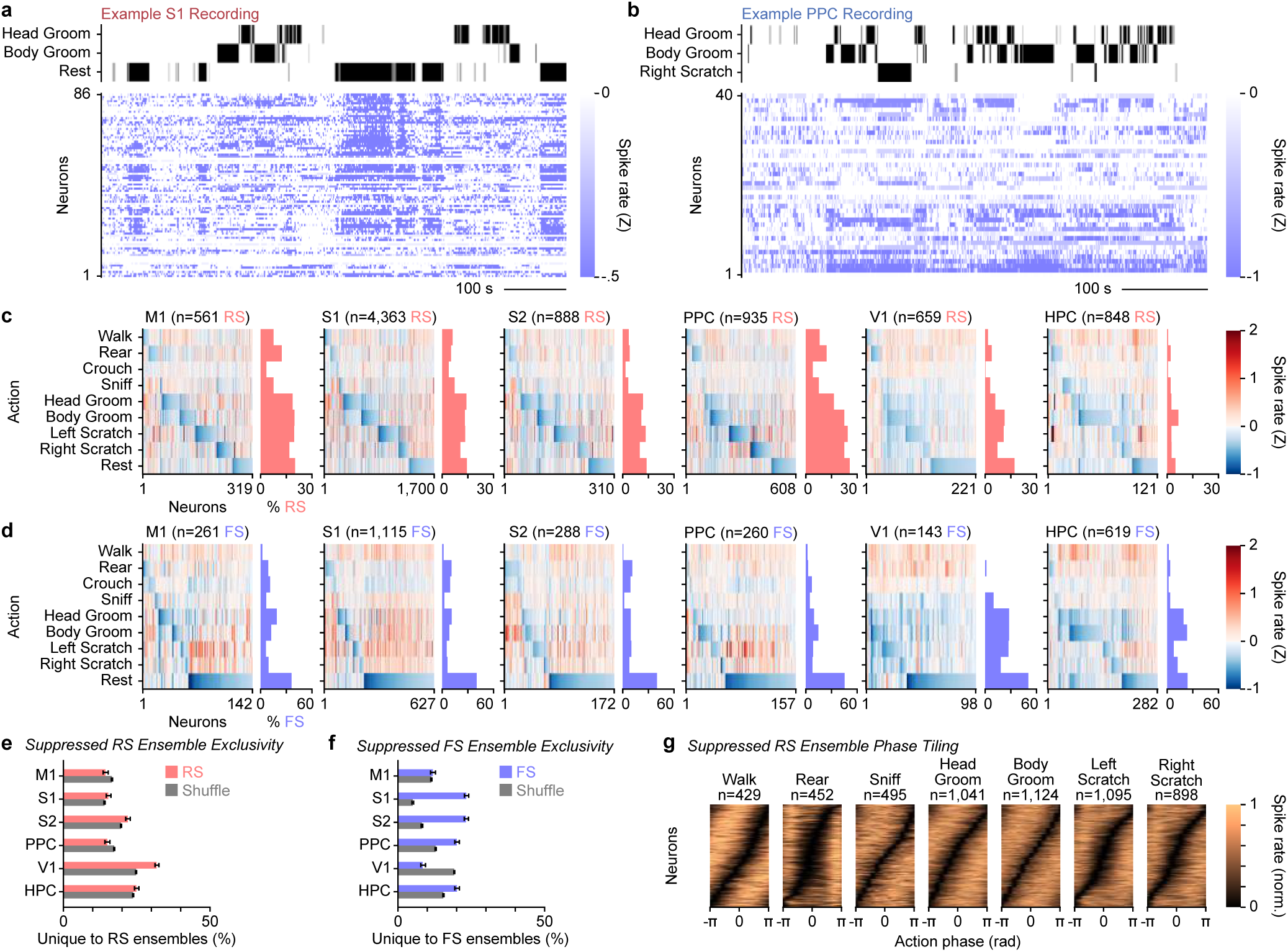
Neural ensembles suppressed during specific actions. **a,** Snippet from behavioral and S1 recordings in an example mouse. Top, Ethogram showing Head Groom, Body Groom, and Rest actions. Bottom, Spike raster aligned in time, sorted by Rastermap. **b**, Same as (a), but for PPC neural activity aligned to Head Groom, Body Groom, and Right Scratch action ethogram. **c**, Heatmaps showing mean Z-scored spike rate of action-suppressed RS neurons (columns) for each action (rows). Histograms showing percentage of all RS neurons belonging to each (non-exclu-sive) suppressed ensemble for each action across cortical regions. **d**, Same as (c), but for FS neurons suppressed during actions. **e-f**, Mean percentage of unique neurons in each suppressed ensemble not shared with other ensembles, compared to unique neurons drawn from shuffled populations (gray) with matched ensemble sizes. Red: RS neurons; blue: FS neurons. Error bars: 95% CI. **g**, Action phase tuning curves for RS neurons in each suppressed action ensemble concatenated across cortical regions, normalized to range 0-1 and sorted by preferred phase.

**Extended Data Figure 7.**
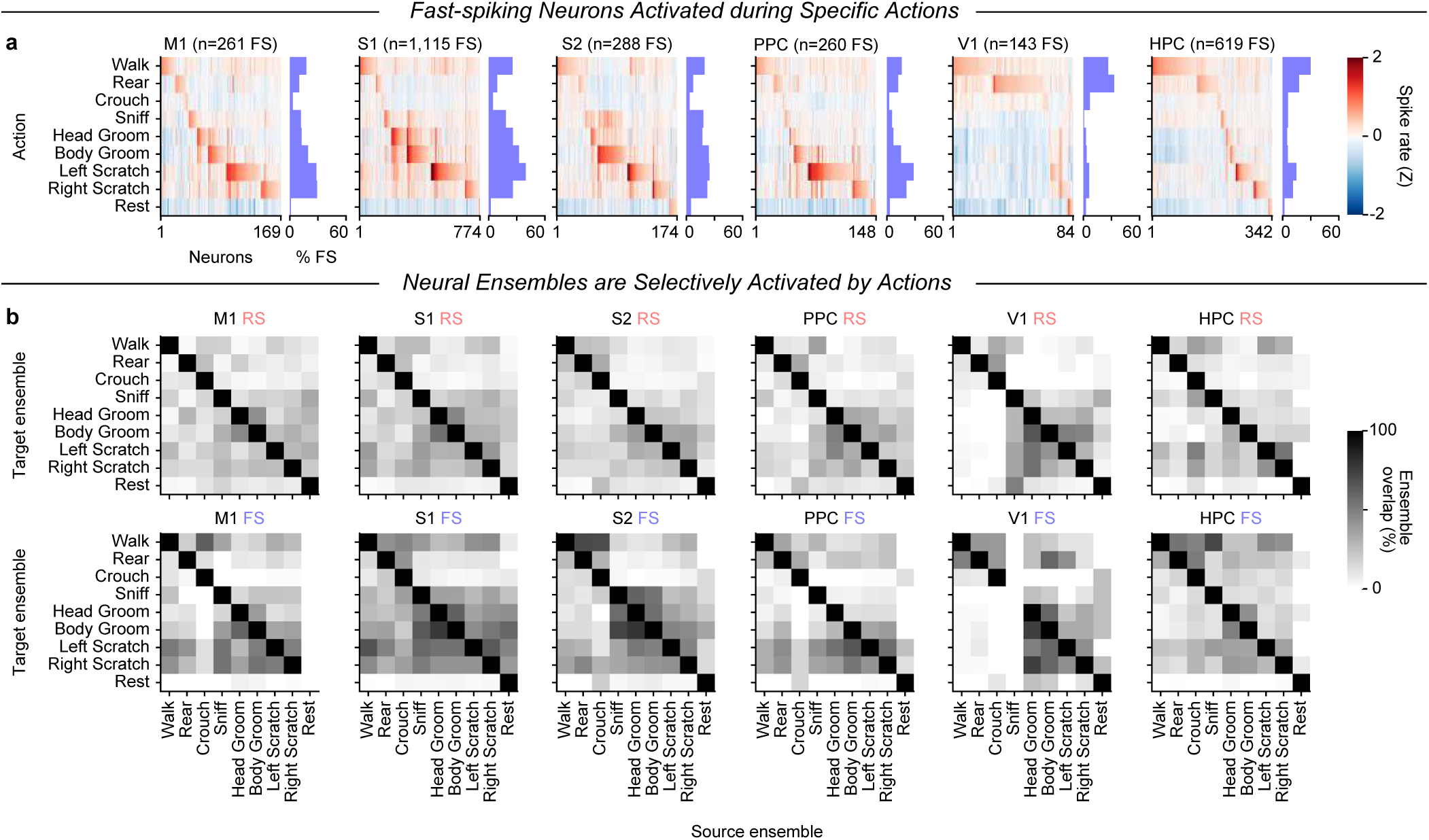
Neural ensembles are selectively activated by specific actions. **a,** Heatmaps showing mean Z-scored spike rates of FS neurons (columns) activated during each action (rows). Histograms showing percentage of FS neurons belonging to each action ensemble. **b,** Heatmaps showing overlap between action ensembles, quantified as the percentage of neurons in each source ensemble (columns) also present in each target ensemble (rows), shown separately for RS neurons (top) and FS neurons (bottom).

**Extended Data Figure 8.**
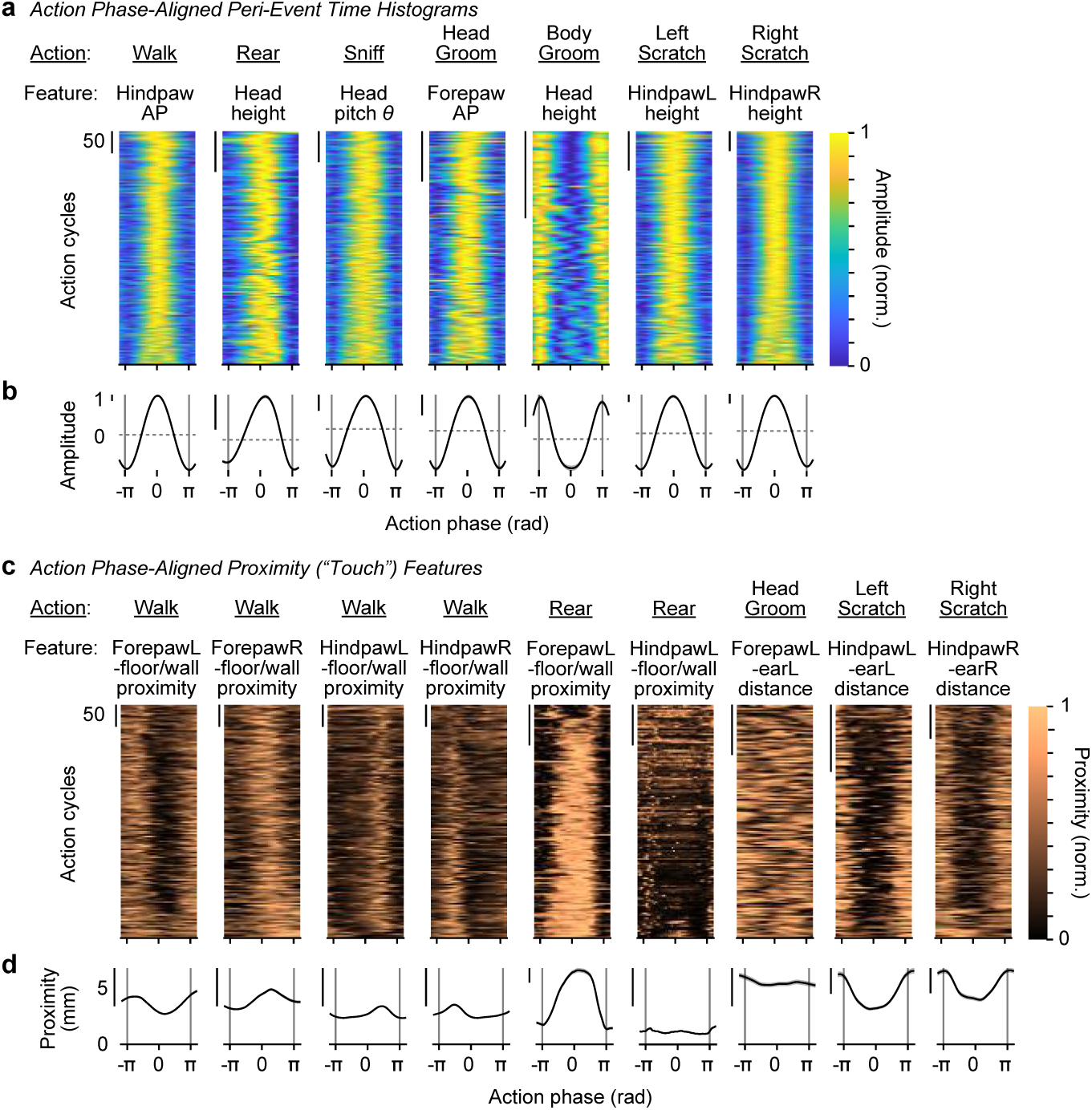
Kinematics and touch-related features are aligned to action phase. **a,** Heatmaps showing action phase-aligned amplitude of kinematic features, normalized for each action cycle (rows). These kinematic features were the ones used to define the phase of each action. Action cycles are concatenated from an example recording for each action. Kinematic features were filtered prior to time-warping to duration of each phase cycle. **b,** Mean phase-aligned amplitude of each kinematic feature, averaged across action cycles. Error shading: 95% CI. Scale bars: 1 mm, except: Rear, 1 cm; Sniff, 1°. **c,** Heatmaps showing action phase-aligned proximity features (listed above), used as a proxy to measure touch events, normalized for each action cycle (rows). **d,** Mean phase-aligned proximity (in mm). Error shading: 95% CI. Scale bar: 5 mm.

**Extended Data Figure 9.**
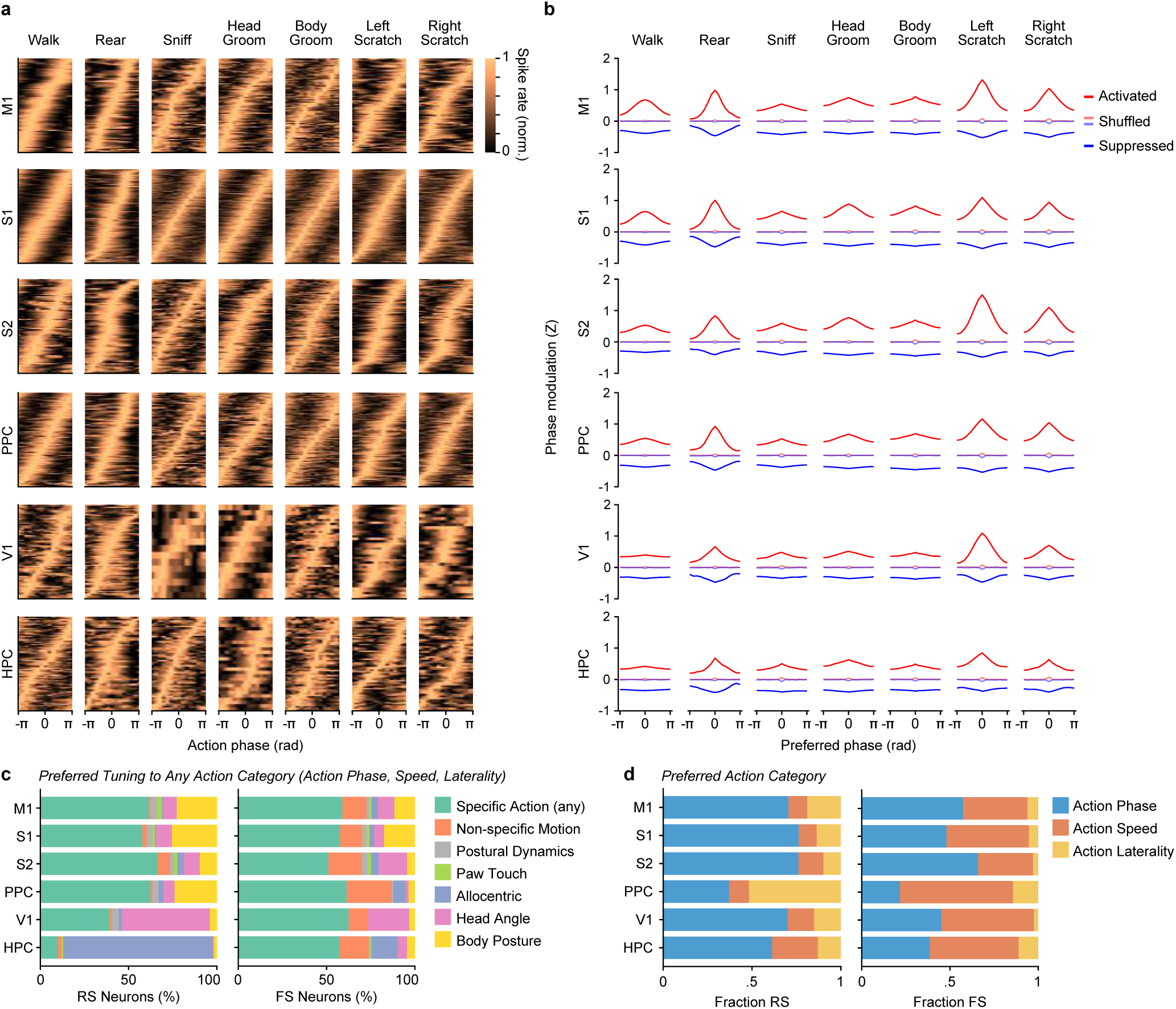
Action-aligned ensemble sequences distributed across cortical regions. **a**, Action phase tuning curves for neurons in each activated action ensemble for each cortical region, normalized to range 0-1 for each neuron (rows). Neuron order within each region is sorted by preferred phase. **b**, Average Z-scored phase tuning, centered on each neuron’s preferred phase, for action-activat-ed (red) and action-suppressed (blue) neurons in each cortical region. Shuffled controls are shown in light red/blue. **c**, Distribution of preferred feature classes for tuned RS (left) and FS (right) neurons across corti-cal regions, with additional action categories (Phase, Speed, and Laterality). Same format as Fig. 5j, but recalculated with Action Phase, Speed, and Laterality in the Specific Actions feature class (see Methods). **d**, Histograms of preferred tuning to Action Phase (blue), Action Speed (red), or Action Laterality (yellow) for RS (left) and FS (right) neurons in each cortical region.

**Extended Data Figure 10.**
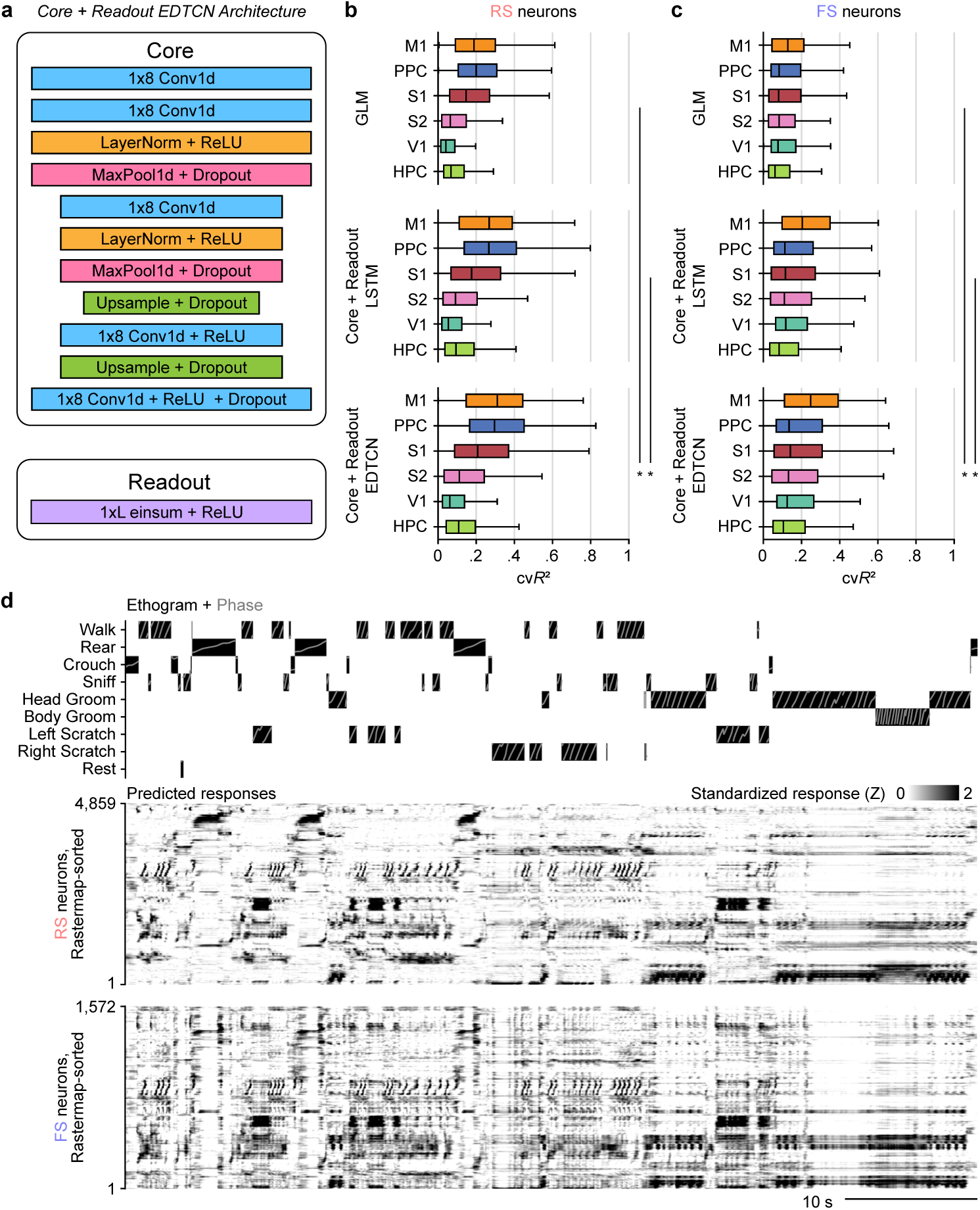
Core + Readout EDTCN outperforms other kinematic encoder models. **a**, EDTCN Core + Readout architecture. The EDTCN Core processes kinematic inputs by compressing the temporal dimension via max pooling and expanding it via upsample layers, producing latent features with the same sequence length as the input. A single-layer Readout module maps these features to predict neural responses for the target frame. **b**, Cross-validated R2 performance of models predicting RS neural activity across cortical areas using generalized linear models (GLM, top), long short-term memory networks (LSTM, middle), and EDTCN (bottom) models. *, Wilcoxon signed-rank test (one-sided), p-val-ues < 1×10^-^¹⁶, corrected by BH-FDR across cell type and model. Median ΔR² EDTCN-GLM, 0.060; EDTCN-LSTM, 0.020 (n = 6115 RS neurons). **c**, Same as (b), but for predicting FS neural activity. *, p-val-ues < 1×10^-^¹⁶. Median ΔR² EDTCN-GLM, 0.051; EDTCN-LSTM, 0.020 (n = 2311 FS neurons). **d**, Top, Ethogram of actions classified by a supervised action segmentation model, overlaid with action phase (gray). Middle, Responses of 4,859 RS neurons across M1, S1, S2, and PPC predicted using the EDTCN Core + Readout model from kinematics in an example session. Note that Actions were omitted from the model inputs. Bottom, Same as middle, but for 1,572 FS neurons across the same cortical regions. Neu-rons are sorted by Rastermap and pooled into superneurons. Predicted responses are Z-score normalized to 0-2 s.d. to emphasize activations.

**Supplementary Table 1.** Mouse cohort and experiment metadata. *Footnote*: For each animal subject, we list Mouse ID, sex, strain, date of birth, session dates (MM/D-D/YYYY; ranges inclusive), experiment type(s), and targeted recording area(s). DANNCE labels indicate mice used to generate multi-camera 3D pose-estimation training labels; Ephys, simultaneous behavior and extracellular recordings; “Action labels,” sessions with human-labeled action classes. Area abbreviations: S1, primary somatosensory cortex; M1, primary motor cortex; S2, secondary somatosensory cortex; PPC, posterior parietal cortex; HPC, hippocampus; V1, primary visual cortex; N/A, no recordings.

| Mouse ID | Sex | Strain | Date of Birth | Recording Dates | Experiments | Recording Location(s) |
| --- | --- | --- | --- | --- | --- | --- |
| KSp003 | M | C57BL/6J | 04/15/2020 | 7/5/20, 7/10/20 | DANNCE labels | N/A |
| KSp004 | F | C57BL/6J | 04/15/2020 | 7/10/20, 7/13/20 | DANNCE labels | N/A |
| KSp005 | M | C57BL/6J | 04/15/2020 | 7/10/20 | DANNCE labels | N/A |
| KSp009 | M | C57BL/6J | 08/03/2021 | 12/05/21-12/15/21 | Ephys, DANNCE labels, Action labels | S1 |
| KSp012 | M | C57BL/6J | 08/17/2021 | 11/27/21-12/21/21 | Ephys, DANNCE labels, Action labels | S1, HPC |
| KSp013 | F | C57BL/6J | 11/16/2021 | 2/07/22-3/03/22 | Ephys, Action labels | M1 |
| KSp014 | F | C57BL/6J | 11/16/2021 | 2/07/22-3/04/22 | Ephys, DANNCE labels, Action labels | M1 |
| KSp015 | F | C57BL/6J | 11/16/2021 | 2/09/22-2/19/22 | Ephys, Action labels | M1 |
| KSp016 | F | C57BL/6J | 11/16/2021 | 2/09/22-3/13/22 | Ephys, DANNCE labels | S1 |
| KSp017 | M | C57BL/6J | 01/04/2022 | 4/04/22-5/02/22 | Ephys, DANNCE labels | PPC, HPC |
| KSp018 | M | C57BL/6J | 01/04/2022 | 4/06/22-5/06/22 | Ephys, DANNCE labels | PPC, HPC |
| KSp019 | F | C57BL/6J | 01/04/2022 | 4/19/22-4/28/22 | Ephys, DANNCE labels | HPC |
| KSp020 | F | C57BL/6J | 01/04/2022 | 4/04/22-5/06/22 | Ephys, DANNCE labels | PPC, HPC |
| KSp021 | F | C57BL/6J | 01/04/2022 | 4/09/22-5/05/22 | Ephys, DANNCE labels | S2 |
| KSp022 | M | C57BL/6J | 01/04/2022 | 4/09/22-5/12/22 | Ephys, DANNCE labels | S2 |
| KSp023 | M | C57BL/6J | 05/16/2023 | 8/03/23-8/30/23 | Ephys | S1 |
| KSp025 | M | C57BL/6J | 05/16/2023 | 8/04/23-9/15/23 | Ephys, Action labels | PPC, HPC |
| KSp026 | F | C57BL/6J | 05/16/2023 | 8/05/23-8/25/23 | Ephys | S1 |
| KSp028 | F | C57BL/6J | 05/16/2023 | 8/05/23-8/29/23 | Ephys | S1 |
| KSp029 | M | C57BL/6J | 08/01/2023 | 12/09/23-12/16/23 | Ephys | V1 |
| KSp030 | F | C57BL/6J | 08/01/2023 | 12/10/23-12/18/23 | Ephys | V1 |

**Supplementary Video 1 | Example DANNCE 3D pose estimation.** Example video (30 seconds of data, 0.5x playback speed). Left, Video from one camera view (out of six) overlaid with reprojected keypoint positions (colored points) and links (colored lines). Right, 3D keypoint positions (white points) and links (colored lines), with position centered to and yaw aligned to the lumbar spine segment.

**Supplementary Videos 2-6 | Example action ensemble videos, kinematics, and rasters.** Representative episodes of Rear (2), Head Groom (3), Body Groom (4), Scratch (5), and Walk (6) from mice implanted in S1, shown with simultaneous video (left) and aligned kinematics and ensemble spiking (right). The red bar marks the current video frame; gray lines indicate action phase onset/offset. Kinematic traces defining each action are plotted above raster plots of action-modulated neurons, sorted by preferred phase. A red arrow denotes the neuron whose spikes are played in the audio. Bottom panels show the automatically classified ethogram.

